# A dynamic normalization model of temporal attention

**DOI:** 10.1101/2019.12.21.886051

**Authors:** Rachel N. Denison, Marisa Carrasco, David J. Heeger

## Abstract

Vision is dynamic, handling a continuously changing stream of input, yet most models of visual attention are static. Here, we develop a dynamic normalization model of visual temporal attention and constrain it with new psychophysical human data. We manipulated temporal attention–the prioritization of visual information at specific points in time–to a sequence of two stimuli separated by a variable time interval. Voluntary temporal attention improved perceptual sensitivity only over a specific interval range. To explain these data, we modeled voluntary and involuntary attentional gain dynamics. Voluntary gain enhancement took the form of a limited resource over short time intervals, which recovered over time. Taken together, our theoretical and experimental results formalize and generalize the idea of limited attentional resources across space at a single moment to limited resources across time at a single location.

## Introduction

The visual system receives continuous, time-varying input, which must be dynamically prioritized according to behavioural goals. However, most data and theory on visual perception and attention have been motivated by a static picture of visual processing, focusing on how we see a single image that is isolated in time. Here we generalized a successful static computational theory of visual and attentional processing into a dynamic model, which we constrained using our recently developed psychophysical protocol and new data on the dynamics of temporal attention.

Our theory is based on the principle of normalization. The normalization model explains contextual modulation in neural populations with divisive suppression ^1, 2^. Normalization appears to be widespread in both basic sensory ^2–6^ and higher-order perceptual and cognitive processing ^7–12^. For this reason, it has been described as a “canonical cortical computation” ^1^.

Several models of attention combine sensory normalization with attentional modulation ^6, 12–19^. In these models, attention changes the sensitivity of neural responses to sensory inputs by modulating the gain of the responses. One such model, developed by Reynolds and Heeger ^12^, proposes that attention modulates neural activity before normalization. This formulation has reconciled ostensibly conflicting electrophysiological and psychophysical findings ^12^ and predicted new results that have been empirically confirmed ^20–22^. However, this leading theory of spatial and feature-based attention is static, with no temporal attention component.

Dynamic normalization models have been developed to account for the time courses of neuronal responses ^5, 10, 23–27^ and dynamic sensory processes like adaptation ^2, 28–32^. But these models have not incorporated attention. It has been noted^17^ that differential shunting equations can be used to implement normalization, as in shunting equation models of spatial attention ^33, 34^.

A major challenge in developing a dynamic normalization model of attention is establishing what the attentional gain dynamics actually are. The behavioural time courses of spatial attention have been characterized: voluntary spatial attention takes 300 ms to be allocated, and involuntary spatial attention peaks at 90-120 ms ^35–38^. But visual attention is not only directed to locations in space; it is also directed to points in time.

Temporal attention is the prioritization of visual information at specific points in time – for example, the moment a behaviourally relevant stimulus will appear ^39^. Even with spatial attention fixed at one location, visual temporal attention can be manipulated using temporal precues to specific time points. Such voluntary, or goal-directed, temporal attention affects perception ^40–44^, neural responses ^44–48^, and microsaccades ^49^. Voluntary temporal attention can lead to both perceptual benefits at attended times and perceptual costs at unattended times, relative to when attention is distributed across time ^41^. But the temporal dynamics of attention that lead to these benefits and costs are unknown. Moreover, there are no existing models of voluntary attention to specific time points.

We define involuntary temporal attention as stimulus-driven attentional dynamics that prioritize specific points in time in a non-goal-directed fashion – for example, an increase in attention following a salient stimulus. Involuntary spatial attention transiently enhances visual processing at a stimulated location, and its underlying mechanisms are at least partially distinct from those underlying voluntary spatial attention ^35, 38^. However, the dynamics of involuntary temporal attention (even when spatial attention is fixed) are unknown, and there are no general-purpose models of involuntary temporal attentional dynamics.

We developed a normalization model of dynamic attention that can capture not only spatial and feature-based attention but also temporal attention. We performed a psychophysical experiment to measure how voluntary and involuntary temporal attention affect perception across time, and we used these new data on temporal attentional dynamics to constrain the model. The model that best fits the data predicts a limitation in the availability of voluntary attentional gain across time intervals of ~1 s. We then used the model, with the same neuronal and attentional parameters, to fit two previous data sets ^41, 42^, thereby providing empirical evidence for the generalizability of our new model.

## Results

### Behaviour

To determine the dynamics of voluntary and involuntary temporal attention, we performed a behavioural experiment (**Figure 1a,b**). Observers judged the orientation of gratings while voluntary temporal attention was directed to different points in time. On each trial, two gratings appeared in sequence at the same location, separated by a stimulus onset asynchrony (SOA). The SOA ranged from 100-800 ms across testing sessions but was fixed within a session to ensure predictable stimulus timing. Voluntary temporal attention was manipulated by an auditory precue to attend the first target (T1), the second target (T2), or both targets (neutral precue). When a single target was precued (80% of all trials), precue validity was 75%: On valid attention trials (60% of all trials), observers were asked at the end of the trial to report the orientation of the precued target; on invalid attention trials (20%), they were asked to report the target that was not precued. On neutral trials (20%), observers were equally likely to be asked to report T1 or T2. Therefore, only the time point(s) to which voluntary attention was directed varied from trial to trial.

**Figure 1.**
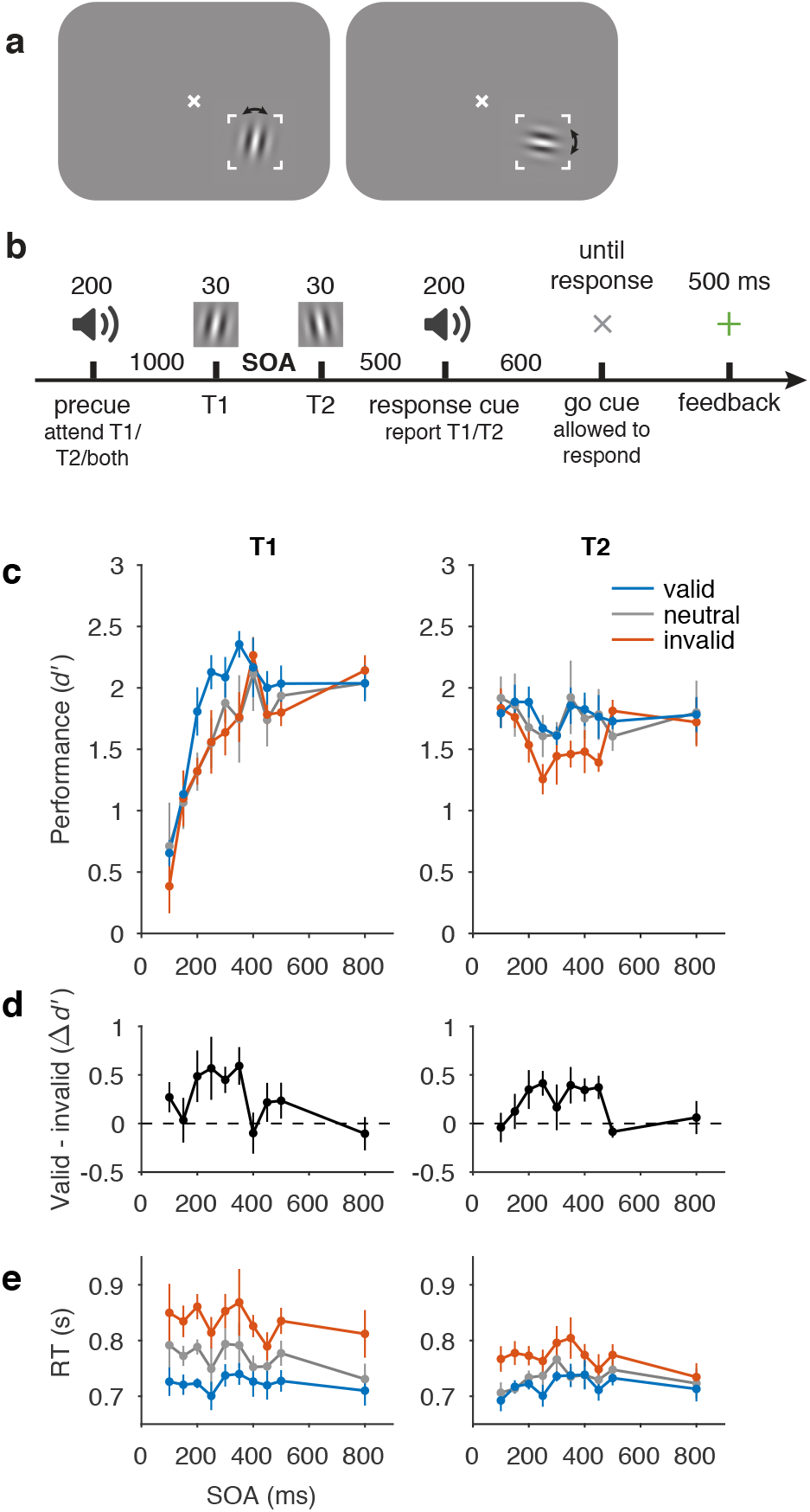
Behavioural protocol and data. **a)** Stimulus display and task schematic. Observers discriminated clockwise (CW) vs. counterclockwise (CCW) tilts about either the vertical (left) or horizontal (right) axis. (Arrows were not shown to observers.) **b)** Trial sequence. Tilts and axes were independent for T1 and T2. SOA varied across testing sessions from 100-800 ms but was fixed within each session. **c)** Perceptual sensitivity (*d’*) for each SOA, precue (valid, neutral, invalid), and target interval (T1, T2). Main effect of precue validity, F(2, 8) = 8.85, p = 0.0094, *η*_G_^2^ = 0.048. **d)** Effect of precueing temporal attention on perceptual sensitivity (i.e., difference in performance between valid and invalid precues). Size of precueing effect depended on SOA, F(9, 36) = 3.13, p = 0.0069, *η*_G_^2^ = 0.20. **e)** Reaction time (RT). Error bars in **c** and **e** are within-observers SEM computed separately for each target to reflect differences among conditions for each target. Error bars in **d** are SEM, *n* = 5.

Critically, this two-target temporal precueing protocol, which we developed in Ref. ^41^, allowed us to measure how voluntary temporal attention affected the perception of both targets as a function of SOA–which was necessary to infer voluntary attentional dynamics. Such measurement could not have been accomplished using previous temporal precueing protocols, which presented only one target per trial, so observers could reorient attention to the second time point if no target appeared at the first. This protocol also allowed us to investigate involuntary attentional dynamics by assessing the impact of involuntary attention elicited by T1 on T2 behaviour, as a function of SOA. Behavioural performance depended on the temporal attentional precue, the SOA, and the reported target. We identified four main features of the behavioural data.

First, voluntary temporal attention affected behaviour, resulting in attentional tradeoffs between the two targets. Overall, perceptual sensitivity (*d’*) was highest for valid trials, lowest for invalid trials, and intermediate for neutral trials (**Figure 1c,d**). In a repeated-measures ANOVA with precue validity, SOA, and target as factors, there was a main effect of validity, F(2, 8) = 8.85, p = 0.0094, *η*_G_^2^ = 0.048. Temporal precueing tended to produce attentional benefits for T1 (valid better than neutral, which was similar to invalid) but attentional costs for T2 (invalid worse than neutral, which was similar to valid). Planned repeated-measures ANOVAs assessing benefits (valid vs. neutral) and costs (invalid vs. neutral) separately for T1 and T2 yielded a marginally significant benefit for T1, F(1, 4) = 5.00, p = 0.089, *η*_G_^2^ = 0.079, but no evidence for a significant cost, F(1, 4) = 0.43, p = 0.55, *η*_G_^2^ = 0.0017. Conversely, there was a significant cost for T2, F(1, 4) = 15.10, p = 0.018, *η*_G_^2^ = 0.065, but no evidence for a significant benefit, F(1, 4) = 0.089, p = 0.78, *η*_G_^2^ = 0.0015. Reaction time showed a similar dependence on the attentional precue, with fastest responses for valid trials, slowest for invalid trials, and intermediate responses for neutral trials (**Figure 1e**; main effect of validity, F(2, 8) = 21.92, p < 0.001, *η*_G_^2^ = 0.27), confirming that speed-accuracy tradeoffs did not drive the differences in *d’*. The presence and pattern of precueing effects indicates temporal attentional tradeoffs across time, consistent with our previous findings with a 250 ms SOA ^41^.

Second, the temporal precue affected perceptual sensitivity differently at different SOAs. The precue had its largest effects at intermediate SOAs, 200-350 ms for T1 and 200-450 ms for T2, and little or no effect at the shortest and longest SOAs. This SOA dependence can be seen in **Figure 1d**, where we plot the difference between *d’* values for trials with valid and invalid precues. Confirming this observation, a repeated measures ANOVA of the precueing effect (valid – invalid) with target and SOA as factors showed a main effect of SOA, F(9, 36) = 3.13, p = 0.0069, *η*_G_^2^ = 0.20. There was neither a main effect of target, F(1, 4) = 0.11, p = 0.76, *η*_G_^2^ = 0.0059, nor an interaction between SOA and target, F(9, 36) = 1.00, p = 0.45, *η*_G_^2^ = 0.10. The pattern of precueing effects was consistent across observers (SI, **Extended Data Figure 1**).

Third, the overall performance of T1 increased substantially with SOA, from *d’* of ~0.6 at the 100 ms SOA to ~2.1 at the 800 ms SOA (**Figure 1c**) on average across precueing conditions, two-tailed paired *t*-test, t(4) = 5.72, p = 0.0046, Cohen’s *d* = 2.56, mean difference and 95% CI = 1.49, [0.77 2.21]. We call this rising function of SOA for T1 “masking-like behaviour” ^50, 51^. The high T1 performance levels for the longest (800 ms) SOA suggests that memory maintenance was not a limiting factor in the performance of this task.

Fourth, the overall performance of T2 exhibited a dip at intermediate SOAs for all precueing conditions, which reached its lowest average point at 250 ms (**Figure 1c**). The dip was larger for invalid trials (reaching *d’* = 1.3 vs. maximum 1.8), but was also present for valid and neutral trials (*d’* = 1.6 vs. maximum 1.9). This U-shaped function of SOA for T2, including its timing, resembles the attentional blink (AB). The AB refers to a difficulty in reporting the second of two targets in a rapid visual sequence, when the targets are 200-500 ms apart ^52, 53^, and it has been much investigated both experimentally and through modeling ^52, 54, 55^. The similarity to the AB includes the so-called “lag-1 sparing,” which refers to the fact that T2 performance is not impaired in AB tasks at short SOAs of ~100 ms ^56^.

Statistically, the variation of *d’* across SOAs and targets was demonstrated by an effect of SOA on *d’*, F(9, 36) = 3.60, p = 0.0028, *η*_G_^2^ = 0.19, which differed for the two targets, SOA x target interaction, F(9, 36) = 15.38, p < 0.001, *η*_G_^2^ = 0.27. For T2 specifically, the *d’* difference between 100 ms and 250 ms, on average across precueing conditions, was also significant, two-tailed paired *t*-test, t(4) = 2.95, p = 0.042, Cohen’s *d* = 1.32, mean difference and 95% CI = 0.34, [0.020 0.65].

For RT (**Figure 1e**), there was a trend toward faster T2 responses than T1 responses, F(1, 4) = 5.52, p = 0.078, *η*_G_^2^ = 0.091, and the precue influenced RT less for T2 than for T1, validity x target interaction, F(2, 8) = 6.83, p = 0.019, *η*_G_^2^ = 0.048. No other main effects or interactions were significant for *d’* or RT, F<1.3 (SI, **Extended Data Table 1**).

To summarize, the psychometric time courses for the two-target temporal precueing task were quite rich, with masking-like behaviour for T1, AB-like behaviour for T2, and the strongest impact of temporal attention on perceptual sensitivity at intermediate SOAs for both targets. These data provide constraints on possible voluntary and involuntary attentional gain dynamics.

### Model

#### General framework

We developed a dynamic perception and attention model in which neural responses are dynamically adjusted through the recurrent processing of a multi-layer neural network. The model describes how perceptual and decision representations evolve over time, through interactions of sensory inputs and attention. The model components are well established in static models of visual cortical function; here we introduced the new dimension of time. Specifically, the model is a generalization of the Reynolds and Heeger (R&H) normalization model of attention ^12^ into the time domain. We call the present model a “normalization model of dynamic attention.”

The model instantiated the hypothesis that the dynamic interactions between attention and orientation perception can be characterized as changes over time in the gain of visual cortical neurons. Gain control is an established mechanism mediating spatial attention ^12, 15^ and has also been implicated in the effects of rhythmic expectation on perceptual sensitivity ^57–59^.

Each layer of the model consisted of a population of neurons whose responses followed the R&H equation (**Figure 2**). Each neuron’s response was determined by the same basic operations: bottom-up input to a neuron in a given layer was filtered through that neuron’s receptive field, multiplied by top-down attentional modulation—which we term “attentional gain”*—*and then divisively normalized by the activity of its neighbors.

**Figure 2.**
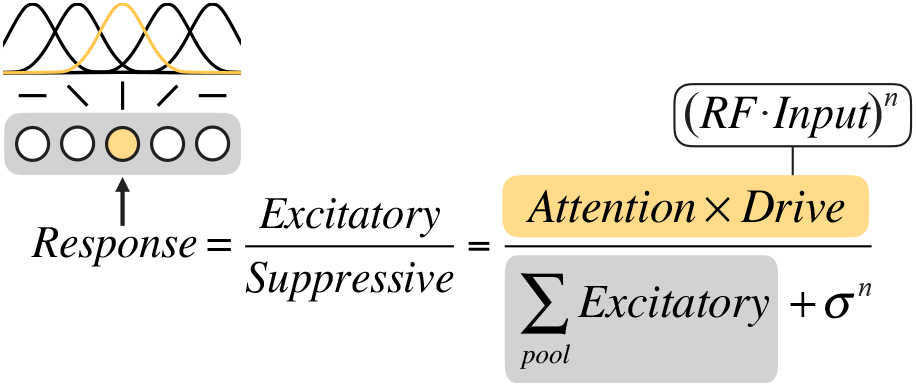
Graphical depiction of the static Reynolds & Heeger normalization model of attention equation. We consider a population of orientation-tuned neurons (illustrated at left, with gray shaded background). The computation of the response of one of the neurons (shaded yellow, with corresponding yellow receptive field (RF)) is shown. The excitatory drive to that neuron is determined by its input drive and attentional modulation (equation text with yellow background). The suppressive drive is determined by the excitatory drives of all the neurons in the local population (equation text with gray background) plus a constant.

To generalize the model from the original static R&H model to a dynamic model, we expressed the model using differential equations that were updated at every time step according to the R&H equation:

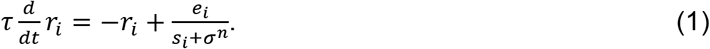

Here *r*_*i*_ is the response of a neuron *i,* where *i* ∈ {1, …, *N*} for a population of *N* neurons; *e*_*i*_ is the excitatory drive to the neuron; *s*_*i*_ is the suppressive drive; *σ* is a semi-saturation constant that keeps the denominator from going to zero and controls the neuron’s contrast gain; *n* is a fixed exponent that also contributes to the shape of the contrast response function; and *τ* is a time constant that determines how long the response takes to rise to steady state when the input turns on and return to zero when it turns off.

The excitatory drive *e*_*i*_ was determined by the equation

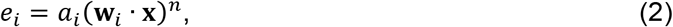

where **x** is the bottom-up input to the layer; **w**_*i*_ is the receptive field (RF) of the neuron; and *a*_*i*_ is top-down attentional gain. Each linear RF computed a weighted sum of its inputs. We describe the inputs **x** and RFs **w** for each layer in Methods.

The suppressive drive *s*_*i*_ was determined by the equation

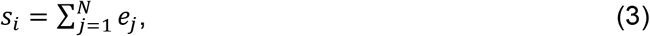

a summation of the excitatory drives of a pool of neurons. Here, the pool was simply all the neurons in the layer (e.g., all orientation preferences at the one spatial location); in general this could be a weighted sum.

At steady state, this differential equation becomes equivalent to the R&H equation; the model therefore retains full generality to predict behavioural and neurophysiological effects of spatial and feature-based attention ^20, 21^, which have been successfully described and predicted by the R&H model ^12^. Like the R&H model, this model is intended to be computationally clear but not biophysically precise; as has been previously discussed ^12^, there are many biophysical mechanisms that could implement normalization. For example, a recently developed circuit model uses recurrence to implement normalization, with steady-state behaviour equivalent to the R&H equation ^27^. In the current model, the “neurons” should be thought of as mapping to computational units at the neural population level.

#### Model specification

The model architecture was a hierarchical, recurrent neural network, with sensory, attention, and decision layers (**Figure 3a**, **Supplementary Table 1**). The layers generated continuous neural response (firing rate) time series given attentional precues and continuous stimulus input (**Figure 3b**). Each layer performed the same computation (Eqs. 1–3); only the inputs and outputs were layer-specific. Full details of the model and simulations can be found in Methods.

**Figure 3.**
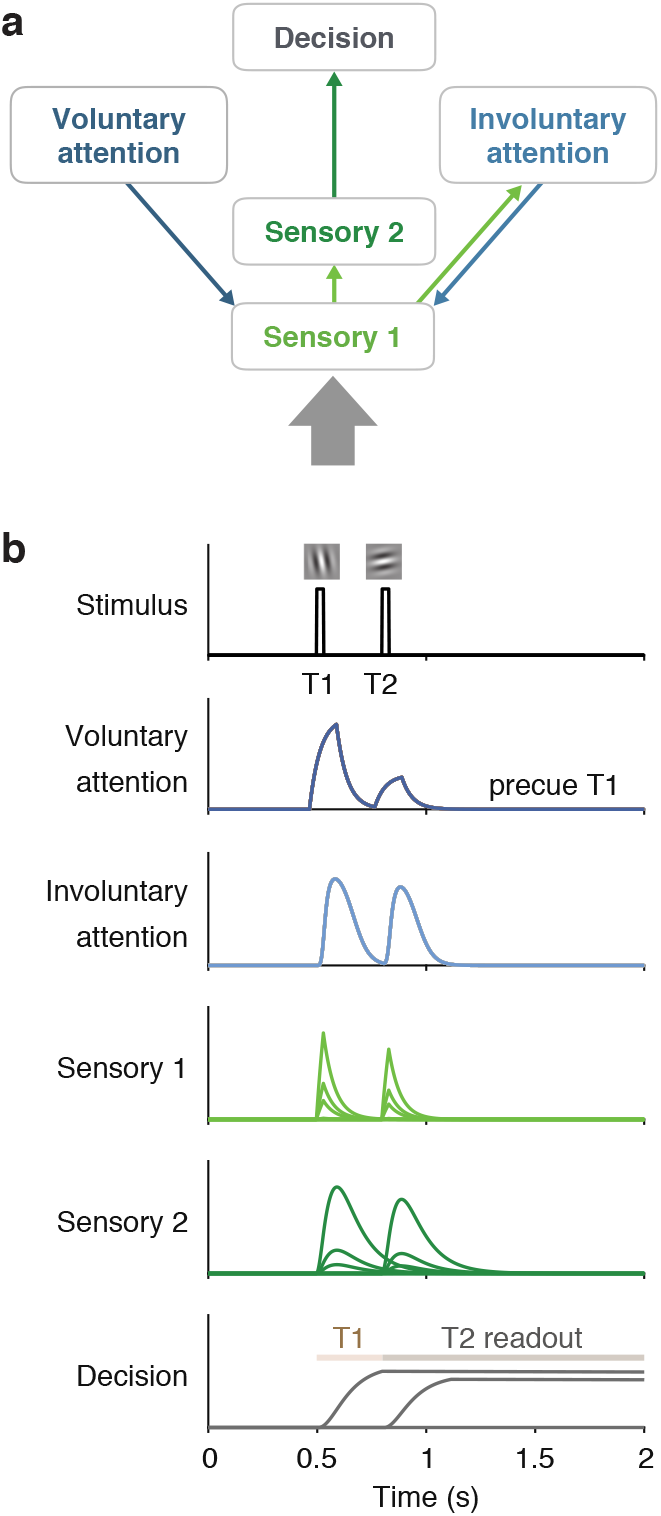
Model architecture and example simulated time series. **a**) Architecture shows sensory input as a thick gray arrow and connections between the layers as thin arrows. An upward arrow indicates input drive, and a downward arrow indicates attentional modulation. **b**) Time series simulated for one trial using a 300 ms SOA, precue to attend to T1, T1 orientation CCW of vertical, and T2 orientation CCW of horizontal, best-fitting parameters. Plots show responses *r* of each neuron in each layer (separate subplots). Separate lines in the sensory layers are neurons with different orientation preferences, and in the decision layer are neurons reading out T1 and T2 responses. The Decision plot shows decision windows as shaded horizontal lines.

##### Sensory layers

The sensory layers represented visual cortical areas. Sensory layer 1 (S1) neurons were orientation selective and received stimulus input. The stimulus orientation (or zero for no stimulus) at every point in time. It also received top-down attentional modulation from both the voluntary and involuntary attention layers. Voluntary and involuntary attention combined multiplicatively to determine the attentional gain *a* for S1. Sensory layer 2 (S2) received input from S1, inheriting its orientation tuning. S2 had a slower rise and more sustained responses than S1 (because the input to S2 was the output from S1), which helped capture T1 behavioural performance as a function of SOA.

##### Voluntary attention layer

The voluntary attention layer (VA) increased attentional gain at task-relevant times. Responses in VA depended on the precue (T1, T2, or neutral) and the trial timing. The input to VA was a time-varying control signal (**Figure 4**) that reflected the observer’s knowledge of the precue and SOA. The control signal consisted of square wave pulses around the times of each target. Pulse latency and duration were free parameters. The amplitude of each pulse was determined by the allocation of voluntary attention to each target – i.e., more voluntary attention at a certain time point generated a larger pulse at that time. These control pulses in turn determined the VA response and corresponding attentional gain modulation of S1.

**Figure 4.**
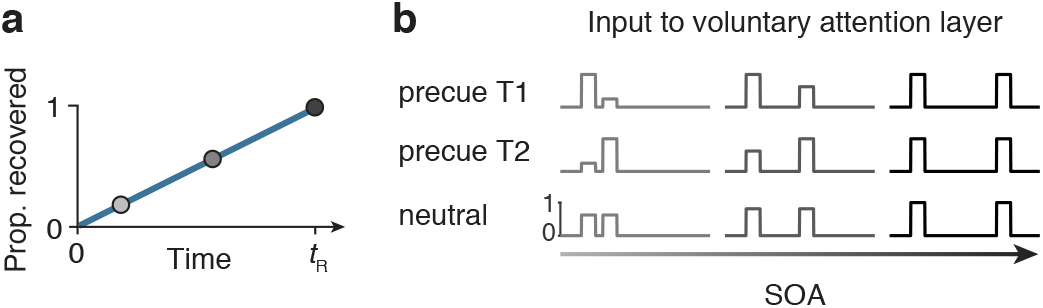
Voluntary attention as a limited but recoverable resource. **a**) Schematic showing the recovery of the input drive to VA, which determined the amplitude of voluntary attentional gain. If maximum voluntary attention was allocated at one moment in time, none was available at the next moment. The available attention then recovered linearly with time, with full recovery at time *t*_R_. **b**) Voluntary attentional control input (*y*) for different precue types and SOAs. When the SOA was short, large attentional tradeoffs between targets occurred due to the limited availability of voluntary attention. When the SOA was long, attention could be allocated maximally to both targets. Voluntary attentional control input was timed, via fitted parameters, to overlap with the sensory responses to each target. In this schematic, equal attention was allocated to T1 and T2 in the neutral condition.

Voluntary attention was a limited resource across time, generalizing the idea of limited spatial attention resources (e.g., Refs. ^21, 35, 60–63^) to the time domain (see “Modeling the data”). Immediately after a maximum (=1) allocation of attention, none was available, but over time attention recovered (**Figure 4a**). We modeled the recovery of attention as a linear function of time, with the recovery time given by the parameter *t*_R_. The precue determined the allocation of attention (**Figure 4b**). When the precue was to T1 or T2, maximum attention was allocated to that target, and as much as possible – given the recovery dynamics – was allocated to the other target. When the precue was neutral, a weighting parameter governed the attentional allocation.

##### Involuntary attention layer

The involuntary attention layer (IA) was stimulus-driven, receiving input from S1. It also fed back to S1, providing a second source of attentional modulation. Because IA responses were driven by S1, they started slightly later than S1 responses (**Figure 3b**). Further, their magnitude depended on the voluntary attentional modulation of S1, because larger S1 responses drove larger IA responses (**Figure 3b**).

##### Decision layer

The decision layer (D) represented a decision area (e.g., in parietal cortex ^64^) and received input from S2. An optimal linear classifier was used to decode CW vs. CCW evidence at each time step from the S2 population. This decoded sensory evidence was the input drive to D. The time constant for D was fixed to be long, which allowed D to accumulate sensory evidence over time, without leak, similar to drift diffusion models ^65^. Decision neurons were target-specific, accumulating evidence only during a corresponding target readout window (**Figure 3b**). The model’s task performance was determined by the response of the decision neuron representing the target that was cued (by the response cue) at the end of the trial.

### Modeling the data

#### Main model

The normalization model of dynamic attention fit the data well (R^2^=0.90) and captured the four main features of the data: (1) voluntary attentional tradeoffs between T1 and T2, (2) largest precueing effects at intermediate SOAs, (3) masking-like behaviour for T1, (4) AB-like behaviour for T2 (**Figure 5a**). Fitted parameter values are listed in **Table 1**.

**Figure 5.**
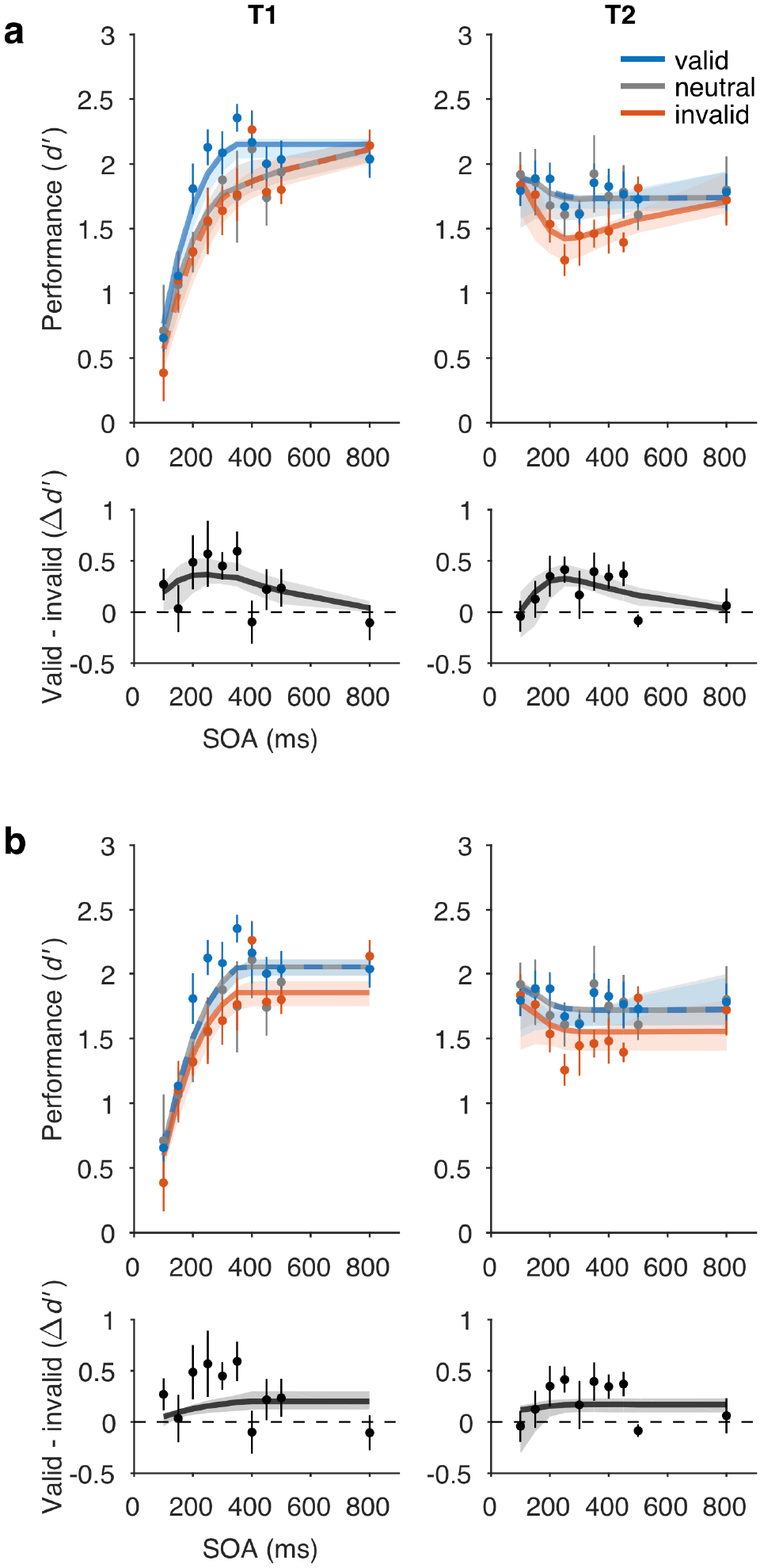
Model fits to perceptual sensitivity data. **a)** Main model with a voluntary attention limit, R^2^=0.90. Top row, performance (*d’*). Bottom row, precueing effect. **b)** No limit model variant, R^2^=0.83. Main model fit better than no limit model variant, ΔAIC=26. Data points and error bars, behavioural performance (copied from Fig. 1, *n* = 5). Curved lines, model fits. Some curves are dashed to reveal overlapping model predictions. Shaded regions, bootstrapped 95% CI for model fits.

**Table 1.**
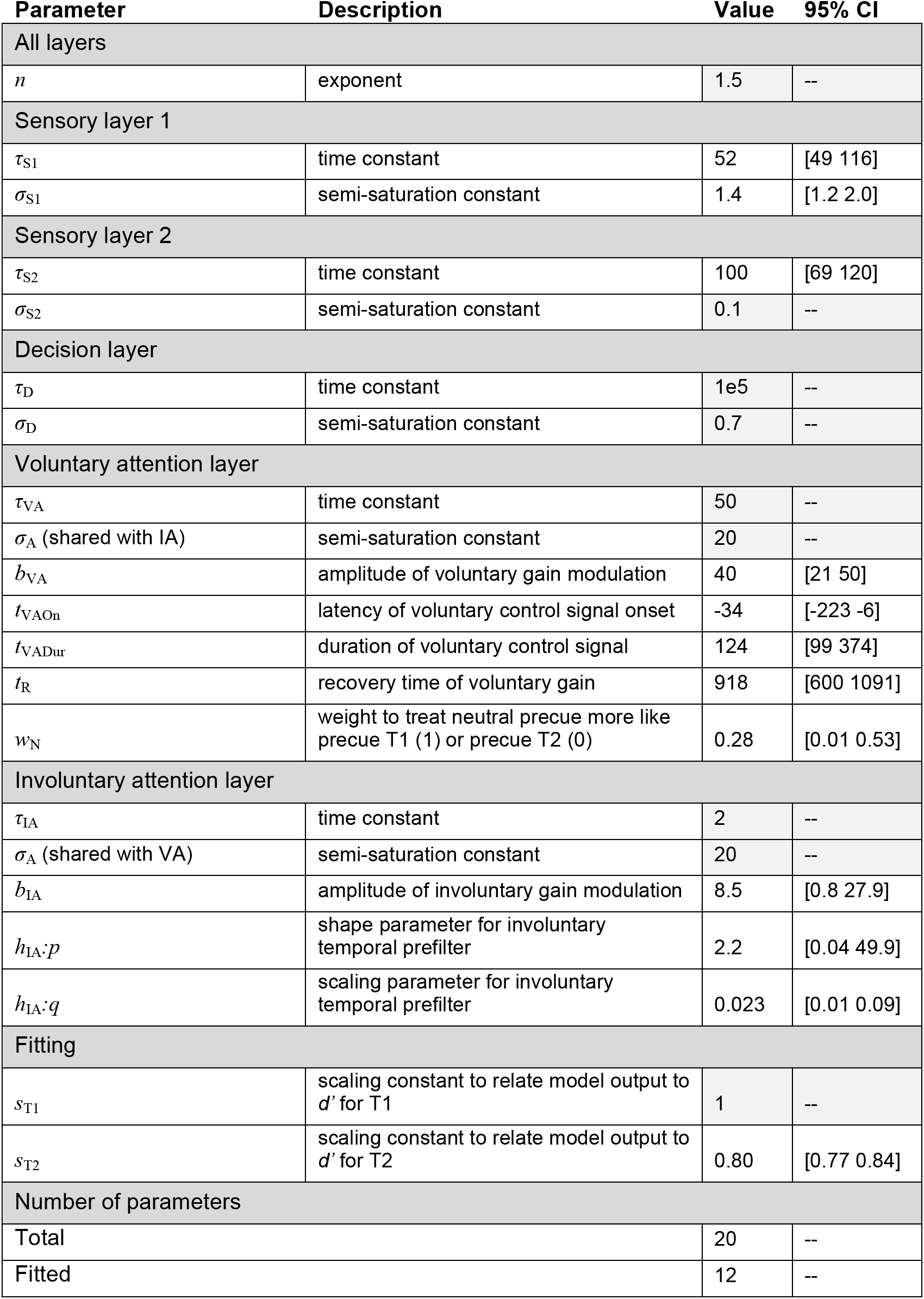
Model parameters. Light gray shading indicates that the parameter was fixed to a set value and not optimized during fitting. All times (i.e., time constants, latencies) are given in ms. Negative latencies for *t*_VAOn_ mean that the voluntary control signal started before the target.

| Parameter | Description | Value | 95% CI |
| --- | --- | --- | --- |
| All layers |  |  |  |
| $n$ | exponent | 1.5 | -- |
| Sensory layer 1 |  |  |  |
| $\tau_{S1}$ | time constant | 52 | [49 116] |
| $\sigma_{S1}$ | semi-saturation constant | 1.4 | [1.2 2.0] |
| Sensory layer 2 |  |  |  |
| $\tau_{S2}$ | time constant | 100 | [69 120] |
| $\sigma_{S2}$ | semi-saturation constant | 0.1 | -- |
| Decision layer |  |  |  |
| $\tau_D$ | time constant | 1e5 | -- |
| $\sigma_D$ | semi-saturation constant | 0.7 | -- |
| Voluntary attention layer |  |  |  |
| $\tau_{VA}$ | time constant | 50 | -- |
| $\sigma_A$ (shared with IA) | semi-saturation constant | 20 | -- |
| $b_{VA}$ | amplitude of voluntary gain modulation | 40 | [21 50] |
| $t_{VAOn}$ | latency of voluntary control signal onset | -34 | [-223 -6] |
| $t_{VADur}$ | duration of voluntary control signal | 124 | [99 374] |
| $t_R$ | recovery time of voluntary gain | 918 | [600 1091] |
| $w_N$ | weight to treat neutral precue more like precue T1 (1) or precue T2 (0) | 0.28 | [0.01 0.53] |
| Involuntary attention layer |  |  |  |
| $\tau_{IA}$ | time constant | 2 | -- |
| $\sigma_A$ (shared with VA) | semi-saturation constant | 20 | -- |
| $b_{IA}$ | amplitude of involuntary gain modulation | 8.5 | [0.8 27.9] |
| $h_{IA:p}$ | shape parameter for involuntary temporal prefilter | 2.2 | [0.04 49.9] |
| $h_{IA:q}$ | scaling parameter for involuntary temporal prefilter | 0.023 | [0.01 0.09] |
| Fitting |  |  |  |
| $s_{T1}$ | scaling constant to relate model output to $d'$ for T1 | 1 | -- |
| $s_{T2}$ | scaling constant to relate model output to $d'$ for T2 | 0.80 | [0.77 0.84] |
| Number of parameters |  |  |  |
| Total |  | 20 | -- |
| Fitted |  | 12 | -- |

To capture the two behavioural features related to voluntary temporal attention – tradeoffs and peak precueing effects at intermediate SOAs – we found it necessary to limit the availability of voluntary attentional gain over time. Specifically, we let voluntary attentional gain be a limited but recoverable resource (**Figure 4**). This property generalizes an idea that is standard in the spatial domain to the temporal domain. In the spatial domain, attention to one spatial location leads to improved processing at that location but impaired processing at other locations, relative to a neutral condition ^60, 61, 63^. Therefore voluntary spatial attention is considered a limited resource at a single point in time that must be distributed across locations. Analogously, in the temporal domain, if such a resource is completely used up at one time point, it will not be available at the next time point; but over time, it will recover to its maximum level. Therefore, within the recovery window, the resource must be distributed across sequential items, leading to tradeoffs. Here, the “limited resource” is, concretely, the allocation of voluntary attentional gain. The estimated recovery time of voluntary attention *t_R_* was 918 ms. Additional quantification of the attentional gain dynamics exhibited by the fitted model can be found in Supplementary **Tables 2 and 3**.

The overall shapes of the performance functions for T1 and T2 were produced by additional model components. The masking-like behaviour for T1 was produced by stopping the decision readout for T1 when T2 appeared. The AB-like behaviour for T2 was produced by a combination of three factors: (1) Limited voluntary attention resulted in lower performance at shorter SOAs, especially for invalid trials. (2) At the shortest SOAs (~100 ms), voluntary attention to T1 was sustained long enough to enhance both T1 and T2 sensory responses, boosting T2 performance. (3) Involuntary attentional excitation combined with voluntary attention to further boost T2 performance at the shortest SOAs, resulting in equal, high performance levels across precueing conditions. A model variant without the involuntary attention layer fit the data almost as well (R^2^=0.89) and was better in model comparison due to having fewer parameters (ΔAIC=−5.8), although it could not produce AB-like behaviour for T2 valid trials (Supplementary Results).

#### No limit variant

A model without limited voluntary attention (**Figure 5b**) produced a poorer fit (R^2^=0.83, ΔAIC=26 with respect to the main model). It also failed to capture the data qualitatively (**Figure 5b**) in two ways. (1) The no limit variant did not produce tradeoffs in temporal precueing effects. It predicted that neutral performance was equal to valid performance for both T1 and T2, unlike in the data, where T1 neutral performance was similar to invalid performance. (2) The no limit variant did not produce peak precueing effects at intermediate SOAs. Rather, the longest SOAs had maximal precueing effects. These failures of the model are due to its structure and could not be altered by a different choice of parameters. A model recovery analysis confirmed the distinguishability of the no limit variant from the main model variant (**Supplementary Figure 1**).

The performance of the no limit model variant reveals why a limit on voluntary attention was necessary. The fact that neutral performance was very similar to valid performance for both targets shows that, without the limit, the model had no incentives to trade off attention between T1 and T2. That is, maximum attention (*y*=1) could be allocated to both targets on every trial with no performance losses. If more attention to one target had led to worse performance for the other, neutral performance would have been worse than valid performance. Indeed, although we built into this model variant a difference between valid and invalid performance by assuming that the observer would follow the precue to attend to one or both targets, the model would have performed the task better overall if it had ignored the precue and attended to both targets on every trial. In that case, the precue would have had no effect on performance at all, unlike what the data showed.

#### Other model variants

To further investigate the necessity of limited voluntary attentional gain in this theoretical framework, we developed two alternative model variants designed to produce attentional tradeoff incentives without a limit on voluntary attention (**Supplementary Figure 2**, **Supplementary Tables 1 and 2**). One model variant had involuntary attentional inhibition, which suppressed T2 more strongly when T1 was precued. The other variant allowed for mutual normalization of late-stage T1 and T2 responses, such that a stronger T1 response would suppress T2 more. However, when these models were fit to the data, neither of these implementations produced tradeoff profiles. We found that limited voluntary attention was still required to let each of these model variants fit the data (**Supplementary Figure 3**, **Supplementary Table 4**), with *t*_R_ estimates of 809 ms and 924 ms, respectively.

#### Generalization to independent data sets and other tasks

To test the ability of the main model to generalize to independent data sets, we fit the model to data from two previous experiments that used the same voluntary temporal attention task ^41, 42^. To do so, we fixed all parameters to the best-fit values from the current experiment and fit only two free parameters to each data set to scale the overall performance of T1 and T2. Thus, the relative magnitudes of the T1 and T2 attention effects and the tradeoffs between benefits to one target (valid vs. neutral) and costs to the other (invalid vs. neutral) were fixed to the values in **Table 1**. The model fit the new data reasonably well, with R^2^=0.83 for the full data sets from each experiment (**Figure 6a,b**). Fits to separate conditions in Ref. ^42^, in which the stimulus was placed at different visual field locations, had R^2^=0.73-0.95 (**Figure 6c**). The model slightly underestimated the precueing effect size for T1 in Ref. ^41^. However, it correctly predicted the smaller precueing effect for T2 compared to T1 in Ref. ^42^, due to the biased attentional tradeoff between targets on neutral precue trials, controlled by *w*_N_. Thus, with parameters for all neuronal and attentional dynamics fixed, the current model could capture independent data sets.

**Figure 6.**
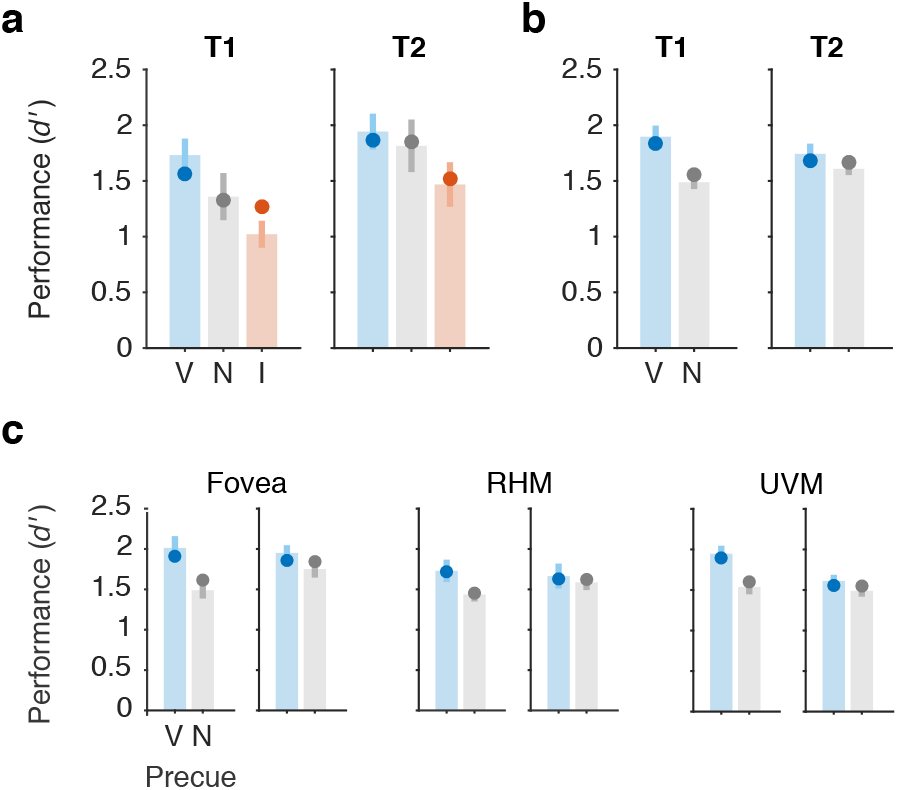
Generalization to independent data sets. The main model was fit (2 free parameters, all others fixed) to previously published data sets: **a)** Denison et al. 2017 Experiment 1, Ref. ^41^, *n* = 10, R^2^=0.83.. **b)** Fernández et al. 2019, Ref. ^42^, *n* = 10, averaged across all conditions, R^2^=0.83, and **c)** separated by visual field location condition, R^2^=0.73-0.95 (RHM = right horizontal meridian, UVM = upper vertical meridian). In both experiments, the two targets were separated by a 250 ms SOA. Bars and error bars show data with SEM. Points show model fits. V = valid, N = neutral, I = invalid.

Finally, we asked whether our modeling framework could capture performance in an AB task. The AB task differs from the two-target temporal cueing task in multiple ways, including (1) targets are embedded in a rapid sequence of non-target stimuli and (2) target timing is unpredictable so voluntary temporal attention cannot be allocated in advance of the targets. A preliminary simulation shows that the current model can capture major features of the AB (**Supplementary Figure 4**) and invites further work testing the normalization model of dynamic attention on the AB and other tasks.

## Discussion

### A normalization model of dynamic attention

We developed a normalization model of dynamic attention, generalizing the Reynolds and Heeger normalization model of attention ^12^ to the time domain. The model is built using components that have support from studies of the visual system and of spatial and feature-based attention, such as linear filters, gain control, rectification, and normalization. Critically, it adds attentional dynamics, i.e., time-varying attentional gain. The model handles temporal attention, including voluntary and involuntary attentional dynamics – in addition to spatial and feature-based attention – in a unified computational framework.

To constrain the model and reveal the dynamics of voluntary temporal attention, we measured how voluntary temporal attention affects perception across time. We found temporal attentional tradeoffs between two sequential targets, which were largest when the targets were separated by SOAs of 200-450 ms. Tradeoffs disappeared at the longest SOAs, revealing a time-limited constraint on processing sequential stimuli that can be accommodated by precisely-timed voluntary control. We also found that the SOA affected the overall performance for the two targets, with masking-like behaviour for T1 and AB-like behaviour for T2.

The model could reproduce the behavioural data using a combination of voluntary and involuntary attentional gain dynamics, together with a simple implementation of masking. Involuntary attention was estimated to be fast and transient, peaking at 82 ms after stimulus onset, consistent with the dynamics of involuntary spatial attention ^38, 63, 66–70^. Although incorporating involuntary attentional gain dynamics into our modeling framework was theoretically motivated, model comparison showed that it was not required to fit the current psychophysical data quantitatively. Future research will be needed to further examine whether and how involuntary attention interacts with voluntary temporal attention.

Voluntary attention took the form of a limited resource that recovered over time, with full recovery estimated to take ~1 s, and consistent with a range of 0.6-1.1 s (95% CI). The attentional limitation in our model could be, for example, either a limitation on the available voluntary attentional gain (***r***^VA^), or a limitation on the activity in voluntary attention control structures (*y*). The model allowed us to separate the dynamics of voluntary attentional gain from other dynamic processes, such as those related to involuntary attention and those leading to masking-like behaviour. It therefore makes specific predictions about different types of attentional gain dynamics (voluntary and involuntary attention time courses). Alternative model variants also required a limitation on voluntary attention across time, but they predicted different gain dynamics. These competing hypotheses could be tested in neurophysiological experiments. The notion of a limited neural resource that can be flexibly allocated is central to multiple domains in psychology and neuroscience, including voluntary spatial attention ^35, 60, 61^ and working memory ^71^. Here we propose a limited resource across time that underlies the selectivity of voluntary attention to points in time.

### Relation to other attention models concerned with dynamics

Previous modeling frameworks that incorporate both attention and some dynamic element include: the “attention gating model”^72, 73 72–74^; the “theory of visual attention” (TVA) ^75–77^; and the “competitive interaction theory” ^17, 33, 34^. Each framework includes different model variants, some of which incorporate normalization ^17, 75^. These models have had success in accounting for behavioural data from various perceptual tasks. Other frameworks focus on rhythmic attention ^78, 79^, which we do not consider here.

There are several important differences between these models and our dynamic attention model. First, we model voluntary temporal attention. TVA has been adapted to model a constant level of expectation across time ^80, 81^, but not attention to specific time points. Second, our model distinguishes between voluntary and involuntary attention, a distinction that is supported by the spatial attention literature ^35, 82, 83^ and has been reported for temporal attention ^78, 84–86^. Third, in previous models ^17, 77^, the role of attention is to control the encoding of sensory signals into working memory. This view of attention differs from our current model, in which attention modulates sensory signals but has no direct role in working-memory encoding. Fourth, our model is built to handle time-varying stimuli and time-varying attention, rather than single, brief displays ^33^, and without being constrained by attentional episodes ^73^. Fifth, our model is explicitly a neural model, built from standard components from visual neuroscience. As such, it makes predictions about the time courses of neural activity that can be tested physiologically.

### Application to the attentional blink?

T2 performance in our two-target temporal precueing task resembled T2 performance in AB tasks ^52^. The fact that we observed AB-like behaviour in a task with no temporal uncertainty, no distractors or masks, and no dual task conditions could help isolate the mechanisms that lead to AB-like behaviour ^87, 88^. The few AB studies in which voluntary temporal attention has been manipulated have reported inconsistent findings ^89–91^. Here, we manipulated voluntary temporal attention and tested different model variants in which voluntary attentional dynamics either contributed to or were independent from AB-like behaviour. In our main model, the attentional blink arises from limited voluntary attention. We found no need to invoke other processes previously proposed to contribute to the AB (e.g., working memory limitations, loss of top-down control, alpha oscillations) ^54, 92, 93^ to explain the AB-like behaviour in our task. However, the contribution of such processes is not excluded by our model, and as yet we have no evidence that our model should be preferred over others to explain the AB per se.

In an influential AB model ^55^, attention is enhanced by the appearance of a target and suppressed during working memory encoding, which leads to the attentional blink. The initial enhancement of attention is similar to involuntary attentional enhancement in our model but the subsequent suppression differs. A neurophysiological AB model proposes that the AB results from a refractory period in the release of norepinephrine by the locus coeruleus (LC), which limits norepinephrine-driven gain enhancement across time ^94^. Future work should examine how voluntary temporal attention affects LC activity; so far there is no evidence that pupil responses, which are influenced by LC, depend on voluntary temporal attention ^95^. As the goal of the current study was to investigate voluntary temporal attention and not the AB, future work will be required to compare alternative models on a variety of tasks in which dynamic attention has been implicated, including the AB task. As a first step, we simulated an AB task and found that our model captures the major features of the AB.

### Future extensions of the model

The current model is a general description of the dynamic interactions between attention and sensory responses. We have focused on how attention affects sensory processing of oriented gratings, a strategy that has proven productive in studies of spatial attention ^12, 15, 21, 35, 61, 63, 68, 82, 83, 96^, facilitated by our knowledge of how orientation is represented in the visual system ^97^. Future work can extend the model to include working memory layers, as sequential processing limitations may also arise in working memory ^54^, as well as more complex feature representations (using additional sensory layers and different RFs) to handle more complex stimuli. It can also investigate how different types of noise at different stages of the model impacts model behaviour. Here the limited resource of voluntary attention was implemented via constraints on the amplitude of the attention control signal over time, resulting in limits on attentional gain. Future work should explore other implementations of a limited resource on voluntary attention over time.

### Conclusion

We developed a model of voluntary and involuntary visual temporal attention, which can serve as a general-purpose computational framework for modeling dynamic attention. Psychophysical measurements revealed perceptual tradeoffs for successive stimuli within sub-second time intervals, which can be controlled by voluntary temporal attention. Precisely timed visual attention may therefore help humans compensate for neural processing limitations over short, behaviourally-relevant timescales. The model predicts that voluntary temporal attentional gain is a limited resource. Future experiments will be needed to test the current model’s predictions and specify the attentional gain dynamics related to spatial and feature-based attention. The time-varying nature of the proposed framework – not to mention of vision itself – calls for new data from psychophysical, neurophysiological, and neuroimaging experiments with dynamic displays.

## Methods

### Behaviour

#### Observers

Five human observers (20-30 years old, 3 female and 2 male) participated in the experiment. All observers provided informed consent, and the University Committee on Activities involving Human Subjects at New York University approved the experimental protocols. Observers were students and researchers at NYU who had experience doing visual psychophysics (though not necessarily temporal attention tasks). All observers had normal or corrected-to-normal vision, and all but author R.N.D. were naïve as to the purpose of the experiment. No statistical methods were used to pre-determine sample sizes, but our sample sizes are similar to those reported in previous psychophysics publications that took a similar approach of collecting a large amount of data per observer (e.g., Ref. ^7^).

#### Setting

The experiment was conducted in a quiet testing room. During experimental blocks, the only light source was the computer monitor. The experimenter was present to give instructions and throughout training and checked on the observer between testing blocks.

#### Stimuli

Stimuli were generated on an Apple iMac using MATLAB and Psychophysics Toolbox ^98–100^ and were displayed on a gamma-corrected Sony Trinitron G520 CRT monitor with a refresh rate of 100 Hz at a viewing distance of 56 cm. Observers’ heads were stabilized by a chin-and-head rest. A central white fixation “x” subtended 0.5° visual angle. Visual target stimuli were 4 cpd sinusoidal gratings with a 2D Gaussian spatial envelope (standard deviation 0.7°), presented in the lower right quadrant of the display centered at 5.7° eccentricity (**Figure 1a**). (The stimulus was placed in this quadrant in anticipation of future neuroimaging studies. We have previously shown the effects of voluntary temporal attention on orientation discrimination to be indistinguishable at different isoeccentric peripheral locations and the fovea, so we expect stimulus location should not impact the results ^42^.) Stimulus contrast was 64%. Placeholders, corners of a 4.25° × 4.25° white square outline (line width 0.08°) centered on the target location, were present throughout the display to minimize spatial uncertainty. The stimuli were presented on a medium gray background (57 cd/m^2^). Auditory precues were high (784 Hz; G5) or low (523 Hz; C5) frequency pure tones, or their combination, and were presented via the computer speakers.

#### Procedure

The task was designed to study the temporal dynamics of voluntary and involuntary temporal attention, including how these two types of attention dynamically interact to affect perception. We used the two-target temporal precueing task developed and previously described by Denison et al. ^41, 49^.

##### Task

Observers discriminated the orientation of grating patches (**Figure 1**). On each trial, two targets (T1 and T2) were presented at the same spatial location, separated by a fixed stimulus-onset asynchrony (SOA, the time interval between the target onsets) on a given day of testing. The target duration was 30 ms. Each target was tilted slightly clockwise (CW) or counterclockwise (CCW) from either the vertical or horizontal axis, with independent tilts and axes for T1 and T2. Tilts ranged from 1.4° to 2.5° across observers. Both horizontal and vertical axes were used to discourage observers from adopting a strategy of comparing the two successive targets on a given trial to judge whether they were the same or different.

##### Overview of experimental manipulations

We manipulated voluntary temporal attention using temporal precues to T1, T2, or both targets. We assumed that the onset of T1 would elicit involuntary temporal attention, and we measured the perceptual effects of involuntary attention on T2 as a function of time by manipulating the SOA. Other time-varying processes that affected the perception of the two targets, like masking, could also be studied as a function of SOA. Because we wanted observers to be able to attend to precise points in time, we eliminated temporal uncertainty by fixing SOA in each testing session, and varied it across sessions. So on a given testing day, the trial timing was constant, and the only thing that varied across trials was the precue to attend to T1, T2, or both targets.

##### Trial sequence

An auditory precue 1000 ms before T1 instructed observers to attend to one of the targets (informative precue, high tone: attend to T1; low tone: attend to T2) or to attend to both targets (neutral precue, both tones simultaneously). Observers were asked to report the orientation of one of the targets, which was indicated by an auditory response cue 500 ms after T2 (high tone: report T1; low tone: report T2). The duration of the precue and response cue tones was 200 ms. For trials with informative precues (80% of all trials), the response cue matched the precued target with a probability of 75% (valid trials) and the other target with a probability of 25% (invalid trials). For neutral trials (20% of all trials), the two targets were indicated by the response cue with equal probability. To reduce the possibility of speed-accuracy tradeoffs, observers were instructed to withhold their response until the fixation cross dimmed (a “go cue”) 600 ms after the response cue. Observers pressed one of two keys to indicate whether the tilt was CW or CCW relative to the main axis, with unlimited time to respond. Long reaction times (> 2 s) were rare, 0.1% of trials. Reaction times were measured relative to the go cue. Observers received feedback at fixation (correct: green “+”; incorrect: red “−”) after each trial, as well as feedback about performance accuracy (percent correct) following each block of trials.

##### Sessions

Three observers completed twenty testing sessions (10 SOAs × 2 sessions/SOA, 9.6K trials total), and two observers completed ten sessions (10 SOAs × 1 session/SOA, 4.8K trials total) on separate days. The SOA order was randomly determined for each observer. Observers who completed 2 sessions/SOA did two sets of 10, with a separate random shuffling for each set. Each session consisted of all combinations of precue type (valid: 60%, invalid: 20%, neutral: 20%), probed target (T1, T2), target tilt (CW, CCW; independent for T1 and T2), and target axis (horizontal, vertical; independent for T1 and T2) in a randomly shuffled order, for a total of 480 trials per session. Observers completed 64 practice trials at the start of each session to familiarize them with the SOA for that day.

##### Training

Observers completed one session of training prior to the experiment to familiarize them with the task and determine their tilt thresholds. Thresholds were found using a 3-down-1-up staircase with all neutral precues at a 250 ms SOA, to achieve an accuracy of ~79% on average across T1 and T2. After determining the tilt threshold, observers completed 64 trials of training with all valid precues, followed by 320 trials identical to an experimental session. The threshold tilt values were used for the remainder of the experiment. (For one observer whose overall performance improved during the first 10 sessions, the tilt was adjusted before the second set of 10 sessions.)

##### Eye tracking

Eye position was recorded using an EyeLink 1000 eye tracker (SR Research) with a sampling rate of 1000 Hz. Raw gaze positions were converted into degrees of visual angle using the 5-point-grid calibration, which was performed at the start of each experimental run. Online eye tracking was used to monitor central fixation throughout the experiment. Initiation of each trial was contingent on fixation, with a 750 ms minimum inter-trial interval. Observers were required to maintain fixation, without blinking, from the onset of the precue until the onset of the response cue. If an observer broke fixation during this period, the trial was stopped and repeated at the end of the run.

#### Statistics

To examine the effects of the experimental manipulations on behaviour, we conducted repeated-measures ANOVAs and calculated *η*_G_^2^ as a measure of effect size using the *ezANOVA* function from the *ez* package in R. ANOVA assumes normality and sphericity. We tested the normality assumption by fitting a linear model to the data and examining the residuals using Q-Q plots and the Shapiro-Wilk normality test in R. The *d’* measure met the normality assumption. The reaction time measure did not; note, however, that reaction time here was a secondary measure that was analyzed to rule out speed-accuracy tradeoffs, ANOVA is typically robust to violations of normality, and the RT results were consistent with the *d’* results. We confirmed that all significant F tests remained significant after Huynh-Feldt sphericity corrections. All statistical tests were two-sided. Cohen’s *d* for paired *t*-tests was calculated as the mean of the paired differences divided by the standard deviation of the paired differences.

### Model

A note on notation: The variables *e*, *s*, *r*, *a*, *g*, *x*, *y*, and *z* are all time-varying, e.g., *e*(*t*), but for simplicity of notation, we omit the time *t*. We use boldface, e.g., **x**, for vectors. As these variables can be both neuron- and layer-specific, we use a subscript to index the neuron and a superscript to refer to the layer, e.g., *e*_*i*_^S1^ for the excitatory drive to the *i*th neuron of layer S1. When no layer superscript is given, we are referring to the general case, applicable to all layers. For constants, which are neither time-varying nor neuron-specific, we use only subscripts.

#### Model specification

The model consisted of a hierarchical, recurrent neural network, with different layers (**Figure 3a**): sensory layers, analogous to visual cortical areas; attention layers, which modulated the sensory responses; and a decision layer, which read out the sensory responses and reported a task decision (here, clockwise vs. counterclockwise grating tilt). All model parameters are listed in **Table 1**.

##### Sensory layer 1

The first sensory layer (**Figure 3b**, S1) represented an early-stage visual area and received stimulus input at every time step. The excitatory drive for each neuron *i* was

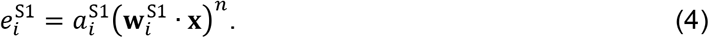

The stimulus **x** was represented in terms of its orientation: a vector of length *M*, with each element corresponding to a different orientation *θ*. When the target grating was on, the element corresponding to the stimulus orientation had value *c*, the stimulus contrast; **x** = (0, 0, …, *c*, …, 0). When the grating was off, all elements were zero-valued. We showed the model the same 2-target trial sequences (**Figure 3b**, Stimulus) as we showed observers (**Figure 1b**).

Layer S1 had 12 orientation-selective model neurons (each of which could represent a larger neural population), which tiled orientation at a single spatial location. Each RF **w**_*i*_^S1^ = (*w_i,1_*, *w_i,2_*, …, *w_i,M_*), was designed to be an orientation tuning curve described by one cycle of a raised cosine,

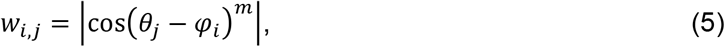

where *θ* is orientation, with sampled orientations indexed by *j*, and *φ_i_* is the preferred orientation of the *i*th neuron. Preferred orientations were evenly spaced, *φ* = (0, *π*/*N*, …, *π*-*π*/*N*). The exponent *m*, which governs the width of the tuning curves, was set to *m* = 2*N*−1. These tuning curves ensure even tiling of orientation space. Orientation selectivity was all that was needed to model our task, but in general the model neurons would be selective also for spatial location and spatial frequency.

S1 received top-down attentional gain modulation from both the voluntary (VA) and involuntary (IA) attention layers, whose responses independently and multiplicatively modulated the sensory drive. The attentional gain *a*^S1^ was

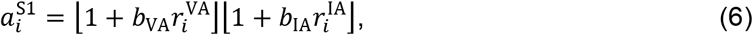

where **r**^VA^ and **r**^IA^ are the responses of the voluntary and involuntary attention layers, respectively; *b*_VA_ and *b*_IA_ are free parameters determining the amplitude of voluntary and involuntary attentional modulation; and brackets denote halfwave rectification, which ensures that attentional modulation is positive. The baseline attentional gain was assumed to be 1, which left sensory responses unchanged in the absence of top-down attentional modulation. When *a* was greater than one, sensory responses increased above baseline; we call this “excitatory” attentional modulation. When *a* was less than one, sensory responses decreased below baseline; we call this “inhibitory” attentional modulation.

##### Sensory layer 2

The second sensory layer (**Figure 3b**, S2) represented a later-stage visual area and received input from S1. Layer S2 also had 12 neurons, and each neuron received input from a single S1 neuron, thereby inheriting the orientation tuning of the S1 neuron (**x**= **r**^S1^). S2 did not receive attentional modulation, so its attentional gain *a* was effectively fixed to 1. The excitatory drive for each neuron *i* was therefore simply

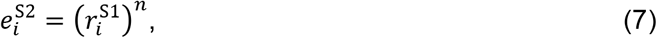

where corresponding neurons *i* in S2 and S1 have the same orientation preference. S2 responses are also determined by equation (1), which includes a temporal low-pass filter with time constant *τ_S2_*. Therefore, S2 had a slower rise and more sustained responses than S1, which helped capture T1 behavioural performance as a function of SOA. However, temporal RFs with more complex dynamics within a layer (e.g., cascade of exponentials) could have achieved the same result using only one sensory layer.

##### Decision layer

The decision layer (**Figure 3b**, Decision) represented a decision area (e.g., in parietal cortex ^64^) and received input from S2. To encode information about temporal order, there were two neurons in the decision layer, one for T1 and one for T2. A binary (0 or 1) decision window *g* gated the input to each neuron: a neuron received input drive only when its decision window was on. The T1 neuron’s decision window started at the onset of T1 and stopped at the onset of T2. The T2 neuron’s decision window started at the onset of T2 and stopped at the end of the trial. Thus, the decision layer read out from successive decision windows for the two targets (**Figure 3b**, Decision, shaded regions). The decision window cutoff for T1 implemented a simplified version of masking, standing in for a mechanism that curtails T1-related signals when T2 appears ^50, 101^.

The input drive to the decision layer was the evidence for clockwise (CW) or counterclockwise (CCW) tilts, based on the responses of S2. The excitatory drive for each neuron *i* was

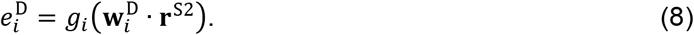

The inner product in this equation represents an optimal linear classifier that decoded CW vs. CCW evidence at each time step from the S2 population response. The classifier projected **r**^S2^ onto the difference **w**^D^ between two templates corresponding to ideal responses to the two possible stimuli, CW and CCW. These templates were “ideal” in the sense that they were the population responses for CW or CCW stimuli at full contrast and with no noise. (We assumed that the orientation axis ‒vertical or horizontal– was known to the observer, so the comparison was only between the CW and CCW templates on the relevant orientation axis.) The classifier projection gave a continuous value that indicated the similarity of the population response on that trial to the CW vs. CCW templates. We arbitrarily assigned CW evidence to positive values and CCW evidence to negative values; an alternative implementation could have used different neurons to represent evidence for each choice. The sign of the evidence indicated a CW vs. CCW choice, and the magnitude indicated the strength of evidence for the choice. Because sensory evidence at each time step created a new input drive to the decision layer, the decision layer accumulated evidence across time, similar to drift diffusion models ^65^. We fixed the time constant of the decision layer to a large value to minimize integration leak and allow faithful evidence accumulation across the trial.

Depending on the response cue, the T1 or T2 decision neuron’s response at the end of the trial was used to determine the model’s performance. Specifically, the model’s CW vs. CCW choice on each trial was determined by the sign of the response, and the magnitude of the decision neuron’s response was presumed to be proportional to the experimentally observed *d’* using a fixed scaling factor. This corresponds to a maximum-likelihood decision assuming additive Gaussian noise. We also fit a separate, relative scaling factor for T2, as the maximum *d’* was lower for T2 than T1. We do not have an explanation for overall differences between T1 and T2, which we have found to vary across datasets and individuals ^41, 42^, but we let these differences be captured by the T2 scaling parameter.

##### Voluntary attention layer

Responses in VA (**Figure 3b**) depended on the precue for a given trial (e.g., precue T1) and knowledge of task timing, so that VA responses increased at the time of the relevant sensory responses. The input *y* to the layer was a time-varying control signal that reflected the observer’s knowledge of the SOA and the precue type ∈ {T1, T2, neutral}. The excitatory drive was determined by the input,

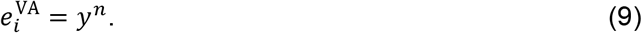

The control signal was a square wave at each target time (**Figure 4**). The timing of each square wave was determined by the SOA and two free parameters, *t*_VAOn_ and *t*_VADur_, which controlled the onset and duration of the square wave. The amplitude of each square wave was determined by the allocation of voluntary attentional gain to each target.

In one model variant, voluntary attentional gain was a limited resource across time. The amplitudes of the square waves then depended on two additional free parameters: a time constant *t*_R_ and an attentional weighting parameter *w*_N_. We implemented the limited resources idea by assuming that immediately after a maximum allocation of gain, no gain would be available, but over time the available gain would recover to the maximum level (**Figure 4**). We modeled the recovery of attention as a linear function of time, with the recovery time given by the parameter *t*_R_. We defined the maximum attention allocation (the amplitude of *y*) at a given time to be 1. Thus the total attention available to both targets for a given SOA was

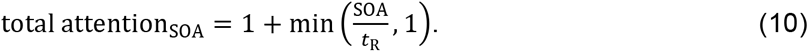

The total attention available across the SOA was distributed across the two targets according to the precue. When the precue was informative (directing attention to one target), the maximum attention for a single target (=1) was allocated to the precued target, and the remainder from the total available attention across both targets was allocated to the other target. (Similar results were obtained when a weighting parameter determined the proportion of attention allocated to the precued target.) For example, if the *t*_R_ was 1000 ms, and the SOA was 400 ms, the total attention available across both targets was 1 (to the precued target) + 400/1000 (to the other target) = 1.4. When the precue was neutral, a weighting parameter *w*_N_ governed the attentional allocation. Due to a perceived or actual asymmetry between the two targets, observers may have a tendency to treat a neutral precue more like a precue for T1 or for T2; *w*_N_ captured this possible bias. **Figure 4** shows examples of the attention control input to VA. We chose the linear recovery function for simplicity and interpretability, as it gives rise to attentional tradeoffs between targets that depend only on *t*_R_ and *w*_N_. Other types of recovery functions, such as exponential recovery, could be explored in future work.

##### Involuntary attention layer

Responses in IA were driven by input from S1 (i.e., “stimulus driven”), as shown in **Figure 3**. The excitatory drive was

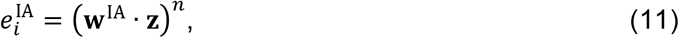

where ***z*** is a temporally filtered version of ***r***^S1^. The temporal filter, which we refer to as a “prefilter”, was a gamma function (*h*_IA_) with amplitude fixed to 1 and fitted shape and scaling parameters *p* and *q*. The gamma function is equivalent to a cascade of exponential (low-pass) filters. The RF **w**^IA^ was the same for all the neurons in the layer and weighted all S1 responses equally, i.e., it was not feature-selective. Because IA responses are driven by S1, they start slightly later than S1 responses (**Figure 3b**). Further, their magnitude depends on the voluntary attentional modulation of S1, with stronger involuntary responses to sensory responses that are voluntarily attended (**Figure 3b**).

#### Model variants

In addition to the main model just described, several variants were developed and tested.

##### No limit model variant

To test whether the limit on voluntary attentional gain was needed to explain the behavioural data, we created a “no limit” alternative. The only difference between the original model variant and the no limit variant was that, in the no limit variant, voluntary attentional gain could be allocated at the maximum level (still defined as 1) at any time, regardless of its allocation at other times. Therefore, this model variant did not require the parameters that determined the allocation of limited voluntary attentional gain in the main model variant: *t*_R_, and *w*_N_.

To give the no limit model variant the best chance to capture the data, we assumed that the model was an “obedient” observer that followed the instruction of the precue to attend to T1, attend to T2, or attend to both targets (neutral precue). So for precue T1 trials, the model allocated voluntary gain maximally (*y*=1) to T1 and minimally (*y*=0) to T2, and vice versa for precue T2 trials. For neutral trials, the model allocated voluntary gain maximally (*y*=1) to both targets. Note that if we had allowed the model to adopt an optimal strategy – that is, to maximize performance accuracy – it would have attended maximally to all targets even for invalid trials, resulting in no difference between valid and invalid trials. This strategy was not available to the variant with limits on voluntary attention.

##### Other model variants

We tested additional variants of the model to assess whether other mechanisms could explain the behavioural data without a temporal limit on voluntary attention. These model variants are described in Supplementary Results.

#### Simulation procedures

The model simulations were run using MATLAB. Each simulated trial lasted 2.1 s with time steps **Δ***t* of 2 ms. The continuous differential equation representing the dynamical version of the R&H normalization equation (1) was discretized as

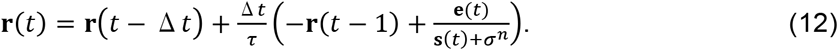

To generate the model performance for one condition (defined by a combination of precue and SOA), a single trial was simulated recursively for 1050 time steps *t*. The only time series specified in advance were the stimulus time series, the voluntary attention control time series *y*, and the decision window time series *g* specifying when evidence would be accumulated for each target. Voluntary attention control was specified based on the precue. The other time-varying quantities – the excitatory drives *e*, suppressive drives *s*, and neuronal responses *r* for each layer – were calculated at each time step, based on the values of the other time-varying quantities at the current and, when prefiltering was applied, previous time steps.

To generate the full psychometric functions containing 60 data points (3 precues × 10 SOAs × 2 targets), each condition was simulated once with no noise to obtain the model performance for both targets in that condition. We performed all simulations with the T1 stimulus CCW from vertical and the T2 stimulus CCW from horizontal, which produced the same behaviour as the average across all possible stimulus sequences.

#### Fitting procedures

We fit each model variant to the group average *d’* data (60 data points). Model fitting was conducted in two phases. In the first phase, the cost function – sum of squared error between model output and data – was evaluated at 2,000 parameter sets sampled from reasonable parameter ranges, which were the same for all models tested. To sample evenly across the full range, each range was divided into 400 bins, and 5 parameter values were sampled uniformly from each bin. In the second phase, the 40 parameter sets with the lowest cost from the first phase were used as starting points for optimization. The optimization algorithm was Bayesian adaptive direct search (BADS) ^102^, which is well-suited for the number of parameters and cost function evaluation time of our model. The optimization producing the lowest cost across all starting points was selected as the best fit.

Some parameters were fixed by hand (i.e., not fit) for theoretical reasons or to minimize redundancies among the fitted parameters. Fixed parameters are shaded in gray in **Table 1**. The values of all fixed and best-fit parameters are given in **Table 1**.

#### Resampling procedures

We obtained confidence intervals on the parameter estimates and model predictions by bootstrapping the data and refitting the model 100 times. For each bootstrap, we aggregated all trials from all observers for each condition, resampled the trials with replacement, and recalculated *d’*. Then, for each resampled dataset, we performed the fitting procedure as described above (2,000 initial cost function evaluations followed by optimization from 40 starting points). This procedure yielded 100 fits of resampled data. Confidence intervals on parameter estimates (**Table 1**) were calculated from the bootstrapped estimates. Confidence intervals on model fits (**Figure 5**) were calculated for each SOA using the bootstrapped model output (predicted *d’*).

#### Parameter interpretation

Several parameters contributed to the attentional dynamics exhibited by each model variant, including the attentional time constants, onset and duration of the voluntary attention control signal, temporal prefilter for the involuntary attention layer, and amplitudes of voluntary and involuntary attentional modulation. We used these fitted parameters to calculate summary metrics describing the voluntary and involuntary attentional dynamics produced by the model. Specifically, we calculated the peak gain amplitudes, the latencies of those peak amplitudes, and the maximum durations of the gain modulations. To calculate the duration of an attentional gain response, we defined a response as non-zero whenever its absolute value was greater than 1% of the maximum response. Note that gain amplitudes are in arbitrary units that should only be compared to the other amplitudes from a given fit.

#### Model comparison

We compared models using the Akaike Information Criterion (AIC), computed with the assumption of a normal error distribution ^103^,

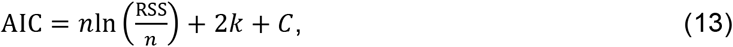

where *n* is the number of observations, RSS is the residual sum of squares, *k* is the number of free parameters, and *C* is a constant that is the same across models. To compare models, we computed the difference in AIC values between two models, ΔAIC. Models with smaller AIC values fit the data better.

#### Generalization to independent data sets

We fit the main model to data from two previous experiments ^41, 42^. Ref. ^41^ Experiment 1 reported discrimination performance as percent correct, so we reanalyzed the data to calculate *d’* from trials aggregated for each precue type. For Ref. ^42^, the data from each visual field location were fit separately, and because there were no significant differences across locations in that study, we also fit the average data across locations. For each fit, two parameters were allowed to vary: *s*_T1_ and *s*_T2_. These parameters are scaling constants that control the overall performance of each target. All other parameters were fixed to the best-fitting values from the main model fit to the current data (**Table 1**). The fitting procedures were otherwise the same as for the current data.

## Data availability

All behavioural data is publicly available on the Open Science Framework (OSF), https://osf.io/dkx7n.

## Code availability

All custom code for the model is publicly available on OSF, https://osf.io/dkx7n. Code for the behavioral experiments is available on GitHub, https://github.com/racheldenison/temporal-attention.

## Acknowledgements

This research was supported by National Institutes of Health National Eye Institute R01 EY019693 to MC and DJH and R01 EY016200 to MC, F32 EY025533 to RND, and T32 EY007136 to NYU. The funders had no role in study design, data collection and analysis, decision to publish or preparation of the manuscript. We thank Hsin-Hung Li for consultation on the model and Carrasco Lab members, especially Valentina Peña for assistance with data collection and Antonio Fernández and Michael Jigo for their comments on the manuscript.

## Author contributions

RND, MC, and DJH conceived of the project, designed the experiment, and interpreted the behavioural results and model findings. RND and DJH conceived of the model. RND implemented the model, conducted the experiment, and analyzed the data. RND wrote and all three authors edited the manuscript.

