## Supplementary Information for "A dynamic normalization model of temporal attention"

### Extended Data

*d'*

| **Effect** | $df_n$ | $df_d$ | $F$ | $p$ | $\eta_G^2$ |
|---|---|---|---|---|---|
| Validity | 2 | 8 | 8.85 | 0.0094 | 0.048 |
| SOA | 9 | 36 | 3.60 | 0.0028 | 0.19 |
| Target | 1 | 4 | 0.011 | 0.92 | 0.00093 |
| Validity x SOA | 18 | 72 | 0.98 | 0.49 | 0.034 |
| Validity x Target | 2 | 8 | 1.29 | 0.33 | 0.010 |
| SOA x Target | 9 | 36 | 15.38 | 6.9E-10 | 0.27 |
| Validity x SOA x Target | 18 | 72 | 0.99 | 0.48 | 0.027 |

Reaction time

| **Effect** | $df_n$ | $df_d$ | $F$ | $p$ | $\eta_G^2$ |
|---|---|---|---|---|---|
| Validity | 2 | 8 | 21.92 | 0.00057 | 0.27 |
| SOA | 9 | 36 | 0.41 | 0.92 | 0.062 |
| Target | 1 | 4 | 5.52 | 0.078 | 0.091 |
| Validity x SOA | 18 | 72 | 0.93 | 0.55 | 0.018 |
| Validity x Target | 2 | 8 | 6.83 | 0.019 | 0.048 |
| SOA x Target | 9 | 36 | 1.23 | 0.31 | 0.016 |
| Validity x SOA x Target | 18 | 72 | 0.73 | 0.77 | 0.0088 |

**Extended Data Table 1.** Repeated measures ANOVA table for behavioral data. SOA = stimulus onset asynchrony, $df_n$ = degrees of freedom in the numerator, $df_d$ = degrees of freedom in the denominator.

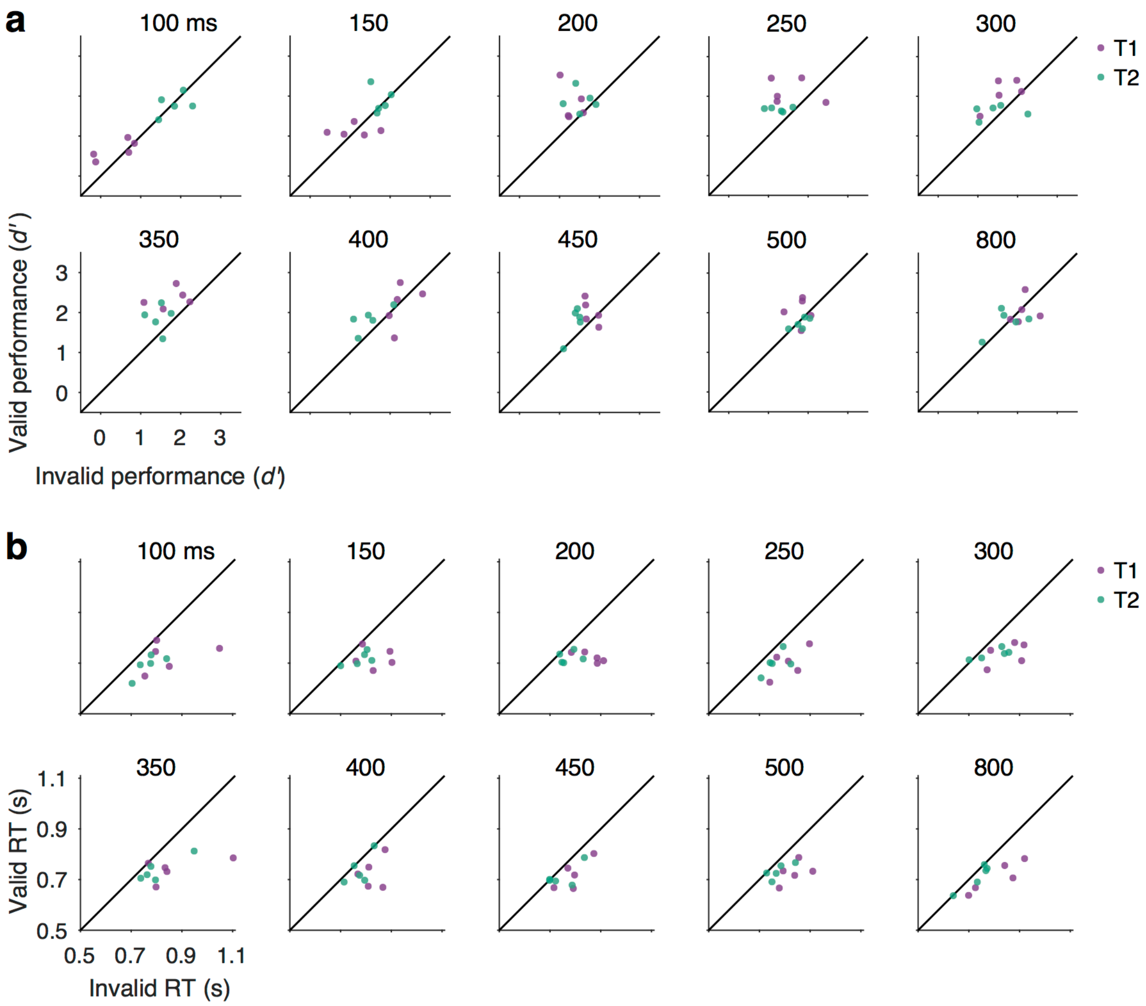


**Extended Data Figure 1.** Behavioral data for individual observers (data points) at each SOA (separate plots). Valid vs. invalid performance for T1 (purple) and T2 (green). For visualization, individual data across SOAs and cuing conditions were normalized separately for each target by adding a constant to equate the individual target mean with the group mean. This adjusts for individual differences in overall performance for a given target without changing the differences among cueing and SOA conditions, facilitating visualization of the pattern of data across these factors. (**a**) Perceptual sensitivity (*d'*). Data points lying above the unity line have a temporal cueing effect: higher *d'* for valid than invalid trials. The improvement of *d'* with temporal attention specifically for intermediate SOAs was consistent across individual observers. (**b**) Reaction time (RT). Data points lying below the unity line have a temporal cueing effect: faster RT for valid than invalid trials. Reaction time improvements were consistent across observers.

### Supplementary Results

#### Model recovery

To confirm the distinguishability of the Main model from its no limit variant, we performed a model recovery analysis. We simulated data from each model variant using its best-fitting parameters. Then we tested the ability of the two model variants to fit these simulated data sets. Each simulated data point was sampled from a normal distribution with mean equal to the model prediction and standard deviation equal to the standard error of the mean across observers. We generated 100 simulated datasets, 50 from each model variant, and then fit all datasets with both model variants. To reduce the computational demand for fitting 100 datasets, we performed optimization using a grid search only (the first stage of our standard fitting procedure). For each dataset, we determined which model variant best fit the data by comparing their AIC scores. The simulated data were more likely to be best fit by the generating model than by the other model, demonstrating successful model recovery (**Supplementary Figure 1**).

Because of the structural differences between the Main and no limit model variants, we expected that errors in model recovery would be driven by the simulated noise. To confirm this, we performed a second model recovery analysis without noise. We simulated one dataset from each model variant using the best-fitting parameters and fit the data using our standard optimization procedure, which includes optimizations from 40 starting points of different parameter sets. For the data generated by the Main model, the Main model always fit better than the no limit model; and vice versa for the data generated by the no limit model variant. That is, the model recovery was perfect in the absence of noise, for all parameter starting points. Together, these analyses show that, in the regime of the empirical data, the models are distinguishable, and the superior performance of the Main model for our empirical data was not merely due to the greater flexibility of that model or some aspect of the fitting procedure.

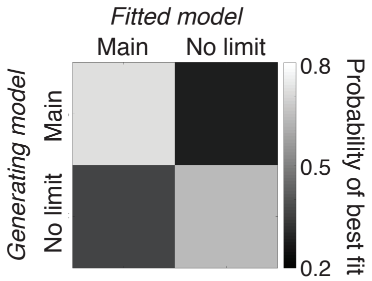


**Supplementary Figure 1.** Model recovery. Probability that each fitted model was the best fit to noisy simulated data generated by each model variant, across 100 simulated datasets.

#### Description of model variants

In addition to the model variant described in the main text (which we call Main), we explored three other model variants that fit the data reasonably well, as well as three counterparts to those models, without a limit on voluntary attention, that were unable to fit the data. Here we describe how each alternative model variant differs from the Main model. **Supplementary Table 1** summarizes the differences among the model variants. A full list of parameters can be found in **Supplementary Table 2**.

| | Sensory layers | Attention layers | Involuntary attention |
|---|---|---|---|
| **Main** | 2 | Voluntary and involuntary | Excitatory |
| **No IA** | 2 | Voluntary | None |
| **EG** | 2 | Voluntary and involuntary | Excitatory and inhibitory |
| **LC** | 3 | Voluntary and involuntary | Excitatory |

**Supplementary Table 1.** Model properties; differences among model variants.

| Parameter | Description | No IA | EG | LC |
|---|---|---|---|---|
| All layers | | | | |
| $n$ | exponent | 1.5 | 1.5 | 1.5 |
| Sensory layer 1 | | | | |
| $\tau_{S1}$ | time constant | 62 | 69 | 47 |
| $\sigma_{S1}$ | semi-saturation constant | 1.4 | 1.4 | 1.3 |
| Sensory layer 2 | | | | |
| $\tau_{S2}$ | time constant | 93 | 83 | 120 |
| $\sigma_{S2}$ | semi-saturation constant | 0.1 | 0.1 | 0.1 |
| Sensory layer 3 | | | | |
| $\tau_{S3}$ | time constant | -- | -- | 2 |
| $\sigma_{S3}$ | semi-saturation constant | -- | -- | 0.3 |
| Decision layer | | | | |
| $\tau_{D}$ | time constant | 1e5 | 1e5 | 1e5 |
| $\sigma_{D}$ | semi-saturation constant | 0.7 | 0.7 | 0.7 |
| Voluntary attention layer | | | | |
| $\tau_{VA}$ | time constant | 50 | 50 | 50 |
| $\sigma_{A}$ (shared with IA) | semi-saturation constant | 20 | 20 | 20 |
| $b_{VA}$ | amplitude of voluntary gain modulation | 48 | 25 | 32 |
| $t_{VAOn}$ | latency of voluntary control signal onset | -28 | -77 | -68 |
| $t_{VADur}$ | duration of voluntary control signal | 152 | 217 | 184 |
| $t_{R}$ | recovery time of voluntary gain | 851 | 809 | 924 |
| $w_{N}$ | weight to treat neutral precue more like precue T1 or precue T2 | 0.34 | 0.24 | 0.20 |
| Involuntary attention layer | | | | |
| $\tau_{IA}$ | time constant | -- | 2 | 2 |
| $\sigma_{A}$ (shared with VA) | semi-saturation constant | -- | 20 | 20 |
| $b_{IA}$ | amplitude of involuntary gain modulation | -- | 5.1 | 19.8 |
| $h_{IA}$:$p_{ex}$ | shape parameter for excitatory component of involuntary temporal prefilter | -- | 1.5 | 2.9 |

| | | | | |
|---|---|---|---|---|
| $h_{IA}$:$q_{ex}$ | scaling parameter for excitatory component of involuntary temporal prefilter | -- | 0.040 | 0.017 |
| $h_{IA}$:$p_{inh}$ | shape parameter for inhibitory component of involuntary temporal prefilter | -- | 20.5 | -- |
| $h_{IA}$:$q_{inh}$ | scaling parameter for inhibitory component of involuntary temporal prefilter | -- | 0.010 | -- |
| $b_{IAinh}$ | amplitude of inhibitory component of involuntary temporal prefilter | -- | 0.48 | -- |
| Fitting | | | | |
| $s_{T1}$ | scaling constant to relate model output to *d'* for T1 | 1 | 1 | 1 |
| $s_{T2}$ | scaling constant to relate model output to *d'* for T2 | 0.82 | 0.83 | 0.81 |
| Number of parameters | | | | |
| Total | | 16 | 23 | 22 |
| Fitted | | 9 | 15 | 12 |

**Supplementary Table 2.** Model parameters. Dashes indicate that the parameter is not used by the model variant. Light gray shading indicates that the parameter was fixed to a set value and not optimized during fitting. All times are in ms.

*No involuntary attention (No-IA).* The No-IA model variant was identical to the Main model variant, but without an involuntary attention layer. The purpose of this model variant was to test whether involuntary attention was necessary to explain the psychophysical data in the present study. **Supplementary Figure 2a** shows the model architecture, and **Supplementary Figure 2b** shows simulated example time series.

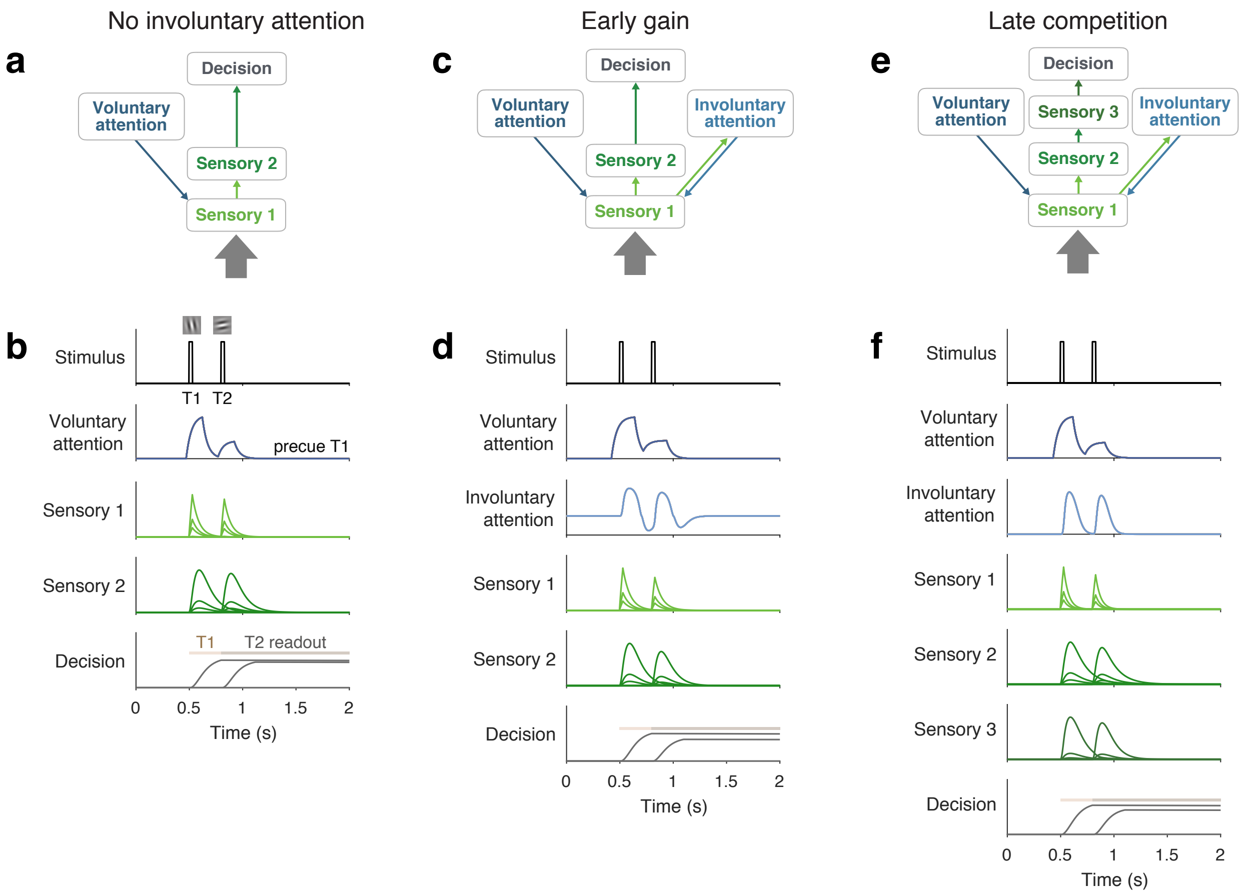


**Supplementary Figure 2.** Model architecture and example simulated time series for each model variant. **a**, **c**, **e**) Architectures show sensory input as a thick gray arrow and connections between the layers as thin arrows. An upward arrow indicates input drive, and a downward arrow indicates attentional modulation. **b**, **d**, **f**) Time series simulated for one trial using a 300 ms SOA, precue to attend to T1, T1 orientation CCW of vertical, and T2 orientation CCW of horizontal. Plots show responses $r_i$ of each neuron (different lines) in each layer (different subplots). The decision layer plots show decision windows as shaded horizontal lines. Time series are scaled to a uniform amplitude for visualization.

*Early gain (EG).* The EG model variant was identical to the Main model variant, except it included inhibition due to involuntary attention. The purpose of this model variant was to test involuntary inhibition as an alternative to limited voluntary attention for creating competition between the two targets. Because this alternative mechanism is a sensory gain modulation, we call this an “early gain” model variant. **Supplementary Figure 2c** shows the model architecture, and **Supplementary Figure 2d** shows simulated example time series.

The involuntary layer of EG had a biphasic temporal response function, which was first excitatory and then inhibitory (**Supplementary Figure 2d**, Involuntary attention). Involuntary attention in the Main model, in contrast, was purely excitatory. To obtain a biphasic temporal response function, we let the temporal prefilter for $\mathbf{z}$ be a difference of gamma functions, which is equivalent to a difference between two cascades of exponential filters. The excitatory and inhibitory gamma functions were governed by shape parameters $p_{\mathrm{ex}}$ and $p_{\mathrm{inh}}$ and scale parameters $q_{\mathrm{ex}}$ and $q_{\mathrm{inh}}$. To control the relative amplitudes of excitation and inhibition, we fixed the excitatory amplitude at 1 and set the amplitude of the negative gamma function to a parameter $b_{\mathrm{IAinh}}$. To allow the neurons to generate both excitatory (positive) and inhibitory (negative) responses while still raising the input drive to the power $n$, we took the difference of a pair of rectified and exponentiated input drives that used temporal filters of opposite phases (one was the negative of the other),

$$e_i^{\mathrm{IA}} = \left\lfloor \mathbf{w}^{\mathrm{IA}} \cdot \mathbf{z}_+ \right\rfloor^n - \left\lfloor \mathbf{w}^{\mathrm{IA}} \cdot \mathbf{z}_- \right\rfloor^n , \tag{1}$$

where the opposite phases are denoted by + and –.

*Late competition (LC).* The LC model variant differed from the Main model variant only in the addition of a third sensory layer (S3). The purpose of this model was to test normalization between sustained neural responses as an alternative to limited voluntary attention for creating competition between the two targets. We call this a "late competition" model variant. **Supplementary Figure 2e** shows the model architecture, and **Supplementary Figure 2f** shows simulated example time series.

The addition of layer S3 allowed normalization over longer timescales. Because normalization applies to a layer's input drive (**Equation 1** of main text), normalization between T1 and T2 responses occur in a given sensory layer only if its T1 and T2 inputs (e.g., responses of the previous sensory layer) overlap in time. In LC, S3 received T1 and T2 inputs from sustained responses in S2, which could result in normalization over longer SOAs compared to a model with only two sensory layers (**Supplementary Figure 2f**).

The excitatory drive of S3 had an identical form to that of S2,

$$e_i^{\mathrm{S3}} = \left( r_i^{\mathrm{S2}} \right)^n . \tag{2}$$

The decision was read out from S3.

Though we have labeled all layers with stimulus representations "sensory" layers, the sustained responses in later sensory layers 2 and 3 could be consistent with either later-stage visual areas like IT or higher-order areas like PFC that are thought to store working memory representations.

*No limit model variants.* To test whether the limit on voluntary attentional gain was needed to explain the behavioral data, we created a "no limit" version of each of the model variants, as was done for the Main model variant.

**Fitting procedures.** The fitting procedures were identical to those described for the Main model variant. **Supplementary Table 2** shows all parameter values (fixed or best-fit) for each model variant. In the LC model variant, the time constant $\tau_{\mathrm{S3}}$ was fixed to a small value, letting S3 inherit the time course of S2, but with the potential for normalization.

**Performance of model variants**

Within the normalization model of dynamic attention framework, we developed three additional model variants. These model variants make different predictions about circuit architecture and attentional gain dynamics. One of the variants tested the necessity of involuntary attention to explain the present data. The other two variants were designed to include mechanisms with the potential for attentional tradeoffs without voluntary gain limits. All the model variants captured the four main behavioral features (voluntary attentional tradeoffs, largest precueing effects at intermediate SOAs, masking-like behavior, and AB-like behavior; **Supplementary Figure 3**) and produced good fits to the data. We found, however, that limited voluntary attention was still required to capture the behavioral data in all these model variants.

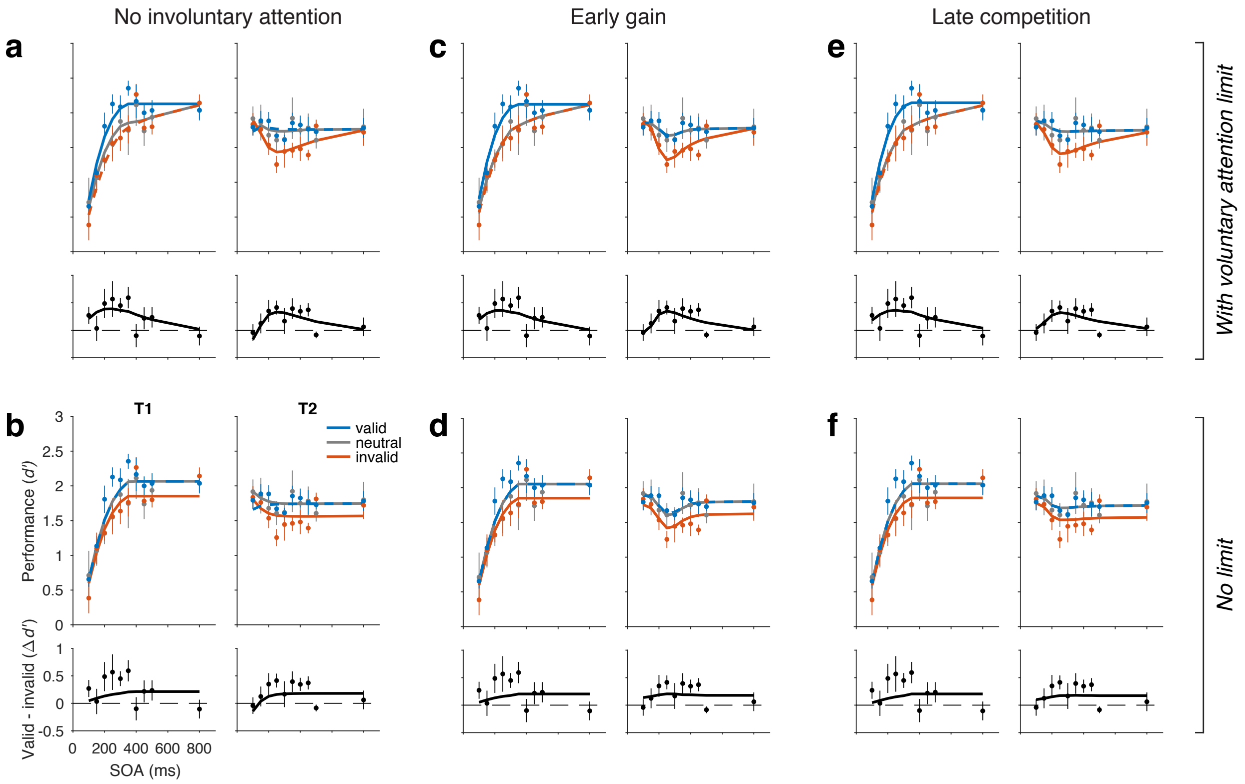


**Supplementary Figure 3.** Model fits to perceptual sensitivity data for each model variant; *d'* and precueing effect data (points) and model predictions (lines). Top row (**a**, **c**, **e**): model variants with a voluntary attention limit. Bottom row (**b**, **d**, **f**): corresponding model variants with no limit. Some curves are dashed to reveal overlapping model predictions. In model variants with voluntary attention limits, neutral overlaps invalid for T1 and valid for T2, similar to the data (tradeoffs). In no limit variants, unlike the data, neutral overlaps valid for both T1 and T2 (no tradeoffs or tradeoff incentives).

**No involuntary attention.** The no involuntary attention (No-IA) model variant performed almost as well, quantitatively, as the Main model variant ($R^2$=0.89) (**Supplementary Figure 3a**). Qualitatively, it was unable to capture any portion of the dip observed around 300 ms for T2 valid and neutral trials. Voluntary attention alone can therefore explain most, if not all, features of the present data. The attentional dynamics of the fitted model are summarized in **Supplementary Table 3**.

*No limit variant.* No-IA-no limit produced a poorer fit ($R^2$=0.82) and had the same qualitative failures as the Main no limit model (**Supplementary Figure 3b**). This result was expected, as there was no additional mechanism in this model to produce tradeoff incentives.

**Early gain.** In the early gain (EG) model variant, involuntary attention had a biphasic (excitatory then inhibitory) temporal response profile (**Supplementary Figure 2c**,**d**), unlike in the Main model variant. The involuntary inhibition contributed to AB-like behavior for T2. EG had potential for tradeoff incentives, because greater voluntary attention to T1 led to stronger sensory responses to T1 and, consequently, stronger involuntary attentional inhibition of T2. This model variant fit the data well ($R^2$=0.90) and captured the four main features of the data

(**Supplementary Figure 3c**). The attentional dynamics of the fitted model are summarized in **Supplementary Table 3**.

*No limit variant.* EG-no limit produced a poorer fit ($R^2$=0.84) and had the same qualitative failures as the Main no limit model (**Supplementary Figure 3d**). Involuntary inhibition did not result in a meaningful difference between valid and neutral performance, i.e., the no limit model did not produce tradeoff incentives.

**Late competition.** In the late competition (LC) model variant, we investigated an alternative mechanism for creating competition between the two targets: normalization between sustained T1 and T2 responses. The only difference from Main was that LC had three sensory layers instead of two (**Supplementary Figure 2e,f**). The third sensory layer was added to allow T1 and T2 responses to normalize one another at longer SOAs (see Supplementary Methods). The normalization between T1 and T2 did not affect T1 performance, because the readout of T1 ended at T2 onset, but it could impair T2 performance. T2 impairment was larger at shorter SOAs, because of greater overlap of the T1 and T2 responses. Voluntary attention to T1 could lead to stronger suppression of T2 responses through normalization, creating potential for tradeoff incentives. This model variant could also fit the behavioral data ($R^2$=0.90). The attentional dynamics are given in **Supplementary Table 3**.

The model did not reproduce the dip in T2 performance around 300 ms for valid and neutral conditions. In principle, the possibility for sustained normalization gave it the capability to do so; however, the fit resulted in relatively short time constants for layer S2, such that T2 performance impairments due to normalization were restricted to SOAs below 200 ms. As a result, limited voluntary attention, and not normalization, was mainly responsible for the AB-like behavior on invalid trials.

*No-limit variant.* LC-no limit produced a poorer fit ($R^2$=0.83) and had the same qualitative failures as the Main no limit model (**Supplementary Figure 3f**). Valid and neutral performance were very similar in the fitted no limit model, indicating no tradeoff incentives, despite normalization.

It is interesting that the EG and LC model variants, when fit to data, entered a regime in which the mechanism designed to generate tradeoff incentives had little impact on performance. Future work will be required to determine whether such tradeoff-inducing mechanisms, when combined with mechanisms for the strategic allocation of temporal attention, would allow parameter solutions that do not depend on limited voluntary attention.

#### Voluntary attention

| | *Excitatory response* | | | | *Recovery* |
|---|---|---|---|---|---|
| | **Peak latency** | **Peak amplitude** | **Normalized amplitude** | **Duration** | $t_R$ |
| **Main** | 88 ms | 1.92 | 1 | 348 ms | 918 ms |
| **No IA** | 122 ms | 2.34 | 1 | 376 ms | 851 ms |
| **EG** | 138 ms | 1.26 | 1 | 442 ms | 809 ms |
| **LC** | 114 ms | 1.63 | 1 | 408 ms | 924 ms |

### Involuntary attention

| | *Excitatory response* | | | | *Inhibitory response* | | | |
|---|---|---|---|---|---|---|---|---|
| | **Peak latency** | **Peak amplitude** | **Normalized amplitude** | **Duration** | **Peak latency** | **Peak amplitude** | **Normalized amplitude** | **Duration** |
| **Main** | 82 ms | 0.51 | 0.27 | 324 ms | - | - | - | - |
| **No IA** | - | - | - | - | - | - | - | - |
| **EG** | 90 ms | 0.28 | 0.22 | 192 ms | 270 ms | -0.15 | -0.12 | 334 ms |
| **LC** | 82 ms | 1.23 | 0.75 | 290 ms | - | - | - | - |

**Supplementary Table 3.** Attentional dynamics. Temporal characteristics of voluntary and involuntary attentional responses derived from the fitted parameters. Latencies are with respect to stimulus onset. Amplitude is in arbitrary units, comparable only within a model, so amplitudes normalized with respect to the voluntary excitatory peak amplitude are also given.

### Model comparison

Model comparison metrics are summarized in **Supplementary Table 4**. All model variants had similar $R^2$ values. No-IA had the lowest AIC score, as it had the fewest parameters, so was the best model by this measure. All the no limit model variants had poorer quantitative fits as well as qualitative failures. Although No-IA had the lowest AIC score, we chose to feature the Main model variant, because existing literature supports a distinction between voluntary and involuntary attention. To make the current modeling framework as general as possible, we therefore include both types of attention layers. Although the present experiment provides specific support only for voluntary gain dynamics, the dynamics of involuntary attention (from the best-fit Main model) are consistent with reported dynamics for transient involuntary spatial attention [1].

| | *Limited voluntary attention* | | *No limit* | |
|---|---|---|---|---|
| | **$R^2$** | **ΔAIC** | **$R^2$** | **ΔAIC** |
| **Main** | 0.90 | 5.8 | 0.83 | 31.9 |
| **No IA** | 0.89 | 0 | 0.82 | 26.2 |
| **EG** | 0.90 | 7.0 | 0.84 | 31.6 |
| **LC** | 0.90 | 6.9 | 0.83 | 33.7 |

**Supplementary Table 4.** Model comparison. ΔAIC is with respect to the best model variant, No-IA with limited voluntary attention.

### Attentional blink simulation

We simulated an attentional blink (AB) experiment using our dynamic normalization modeling framework. To simulate a typical AB task, a stimulus was presented every 100 ms. Two of the stimuli (T1 and T2) were targets, and the rest were distractors, with two distractors before T1 and after T2. We tested ten “lags” between T1 and T2, where lag 1 corresponded to a 100 ms SOA. Targets were oriented stimuli tilted clockwise or counterclockwise about either of the cardinal axes. Distractors were oblique oriented stimuli. The decision layer integrated evidence from the cardinal axes only, allowing the model to differentiate targets from distractors. All stimuli elicited involuntary attention in a bottom-up fashion. Only targets elicited voluntary attention. We assumed that both targets elicited the maximum available voluntary attention at their time of appearance. Due to the limited resource of voluntary attention in the model, T1 elicited more voluntary attention than T2 at shorter SOAs.

We compared the performance of the Main model on the AB simulation to the performance of the Early Gain 1 (EG1) model, which has greater flexibility in its attentional dynamics. Both models captured the major qualitative features of the AB (**Supplementary Figure 4**). T2 performance was high at lag 1, capturing the “lag 1 sparing” effect found in the AB literature [2, 3], dropped at lag 2, and returned to a high level performance at the longest lags.

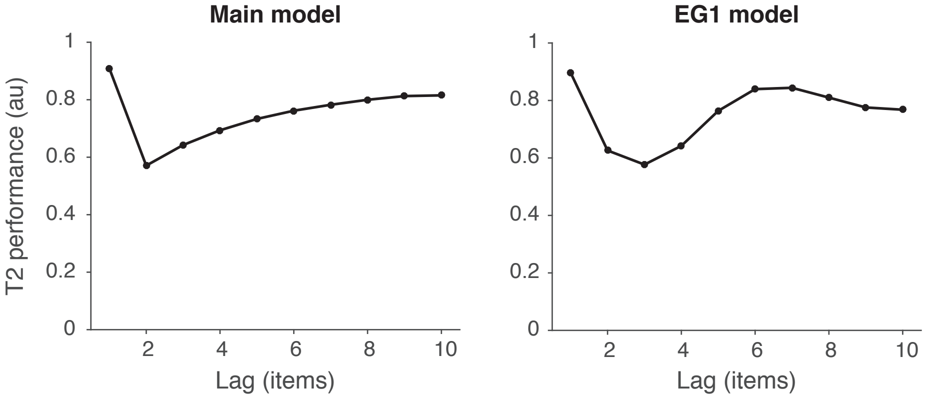


**Supplementary Figure 4.** Attentional blink simulation. The following changes to model parameters were made for the AB simulation: The number of orientation channels was increased to 24 so that oblique distractors would not contribute to cardinal target decision readouts. Because the target times are unpredictable in an AB task, anticipatory voluntary attention is not possible; therefore, the voluntary control signal was set to turn on when the stimulus appeared ($t_{\mathrm{VAOn}}$ = 0 ms). For both Main and EG1, $\tau_{\mathrm{VA}}$ = 20, $b_{\mathrm{VA}}$ = 100, $t_{\mathrm{VADur}}$ = 90. For Main, $\sigma_{\mathrm{S1}}$ = 5. For EG1, $b_{\mathrm{IA}}$ = 20 and $h_{\mathrm{IA}}$:$p_{\mathrm{ex}}$ = 1.

Although the main purpose of the current study was to investigate and model voluntary temporal attention, this simulation shows that the dynamic normalization model of attention framework can generalize to different tasks, such as the AB. The AB protocol differs from the two-target temporal cueing protocol in that it involves: a rapid stream of stimuli, both targets and distractors, unpredictable timing for both targets, no temporal cues, and a requirement to report both targets. In addition, AB studies have generally used letters and numbers as stimuli. In the future, it will be informative to run both the two-target temporal cueing protocol and an AB protocol with Gabors in the same observers and fit the model variants to the combined dataset.
